# Evidence for alternate stable states in an Ecuadorian Andean Cloud Forest

**DOI:** 10.1101/2021.11.26.470126

**Authors:** Ana Mariscal Chávez, Daniel Churchill Thomas, Austin Haffenden, Rocío Manobanda, Laurence Duvauchelle, William Defas, Miguel Angel Chinchero, Danilo Simba, Edison Jaramillo, Bitty A. Roy, Mika Peck

## Abstract

We analyzed a set of historical data from rapid vegetation inventories in a tropical montane cloud forest in northern Andean Ecuador. Trees in plots from several types of forest were counted and measured, including: (1) primary forest, including mature and recently closed-canopy sites and naturally formed gaps, (2) abandoned pasture, and (3) intensively-farmed sites. The goal of the study was to understand in a specific period of time the similarities and differences among natural and anthropogenic disturbances and their potential long term effect on the forest plant community. We found that mature and intermediate close canopy sites are similar. Also old forest is quite resilient to gap-forming disturbance. Natural gaps are quickly colonized by old-forest-associated plant species, and return to an old-forest-type community of trees in a short time. In contrast, forests regenerating from anthropogenic disturbance appear to have multiple possible states: some regenerating forest sites where the anthropogenic disturbance were low are coming to closely resemble old-forest-type communities, but some where the anthropogenic disturbance was intense appear to be changing in a very different direction, which does not resemble any other vegetation community type currently in the forest. A major predictor of present ecological state is the type of land use before reforestation: pastures can occasionally transition back to the pre-disturbance state of forest. More intensively used sites, many of which are abandoned sugar cane plantations, do not return to a pre-disturbance ecological state, instead forming a new and different kind of forest, dominated by a different community of trees. We examined tree-seedling communities to understand the trajectory of the forest into the future, and find that new forest types may be forming that do not resemble any existing associationsintensive agricultural sites. We also found that Los Cedros is extremely diverse in tree species. We estimate approximately 500 species of tree in only the small southeastern area of the reserve that has been explored scientifically. Additionally, the forest tree community shows extremely rapid distance decay (beta-diversity), approaching near complete turn-over in the limited spatial extent of the study. This suggests that hundreds of other tree species remain to be observed in the reserve, in addition to the 350+ that are directly observed in the present study, including new observations of species with IUCN threatened-endangered status. We also highlight the conservation value of Reserva Los Cedros, which has managed to reverse deforestation within its boundaries despite a general trend of extensive deforestation in the surrounding region, and to protect large, contiguous areas of highly-endangered Andean primary cloud forest habitat.

## Introduction

Succession has been a central topic of discussion in ecology for at least a century (Bray & Curtis, 1957; Bunin, 2021; Chambers et al., 2013; Chang & Turner, 2019; Clements, 1916; Connell & Slatyer, 1977; Cowles, 1899; Curtis & Greene, 1949; Gleason, 1926; Holling, 1973; Whittaker, 1956). As with many emergent properties of ecosystems, succession in ecosystems is endlessly controversial: the very existence of stable equilibria states and predictable successional (seral) stages have long been both called-into-question and defended (Bunin, 2021; Gleason, 1926; Norden et al., 2015; Richards, 1952). In recent times increasing attention has been given to models that lie within a community assembly and/or coexistence framework, to explain and predict community composition in plants (HilleRisLambers et al., 2012; T. P. Young et al., 2001). Rather than attempting to broadly predict changes in dominant species associations in the form of seral stages, community-assembly and coexistence models give greater attention to the importance of individuals, which sum to explain differences in community compositions. These models emphasize dispersal, priority, and stochastic effects on individuals, and in the case of niche-based or trait-based models, a further discussion of selection or “filtering” by biotic and abiotic factors (Fisher & Mehta, 2014; Hubbell, 2008; Keddy, 1992; Kraft et al., 2008; Laughlin & Laughlin, 2013; Ulrich et al., 2016). There is no inherent contradiction among these two broad approaches to community modeling: successional and assembly/co-existence approaches are highly complementary (Bhaskar et al., 2014; T. P. Young et al., 2001), and are sometimes packaged into a general “dynamics” framework (Pickett et al., 2009). This cultural shift in community ecology is perhaps due to a desire by ecologists to move away from the highly idiosyncratic, localized knowledge of a site that is often required to successfully predict successional patterns (Norden et al., 2009, 2015), and instead to more abstract and more universal ecological models.

This shift may also perhaps be partly due to the chaotic times in which we find ourselves. In the current era, most ecosystem types are experiencing historical rates of species loss (Barnosky et al., 2011; Pimm et al., 2014), and are already undergoing consequences of climate change (Allen et al., 2010; Carmody et al., 2016; Ciais et al., 2005; Day et al., 2008; Doney et al., 2012; Fadrique et al., 2018; Mantgem et al., 2009). Ecosystems are also continually receiving new species from human activity (Hobbs et al., 2009) and face direct modification or even wholesale elimination due to land use change (Ellis et al., 2010; Hautier et al., 2015; Vitousek et al., 1997). Few anthropogenically disturbed ecosystems seem to have been successfully “returned” to stable historical ecological states (Hobbs et al., 2009), let alone to climax states that resemble some sort of prehistoric ecological stability. In our destabilized era, the search for ancient species associations that require multiple uninterrupted, decadal journeys through intermediate ecological stages can sometimes seem quixotic.

However, the discussion of succession in forests has taken on particular and new urgency in the current era of accelerating climate change. Forests and their soils fix a significant portion of carbon every year, which has probably been essential in slowing negative effects of anthropogenic atmospheric carbon release (Pan et al., 2011). Forests and their soils are also sitting reservoirs of vast amounts of sequestered carbon in soils that are readily lost through disturbance (Lal, 2005). After many decades of underappreciation as essentially carbon-neutral ecosystems (Odum, 2014), a vibrant discussion has arisen around the role of primary forests not only as immense storehouses of carbon, but also as continuing carbon sinks (Gundersen et al., 2021; Luyssaert et al., 2008). New, proforestation-centered prescriptions for climate mitigation have therefore gained support: old forests (primary or late-stage successional) should be be protected as reserves of existing fixed carbon, and potentially high-functioning sinks for further carbon fixation, and management of secondary forests should enhance characteristics that maximize carbon sequestration (Lewis et al., 2019; Moomaw et al., 2019; Sacco et al., 2021; Watson et al., 2018). Such characteristics are often exemplified by old forests (Harmon et al., 1990; Stephenson et al., 2014). Tropical forests are the most volatile portion of the global forest carbon sink, and their role as a sink for atmospheric CO2, rather than a source, will depend greatly on how they are managed and protected in the coming years (Mitchard, 2018). Tropical montane forests are likely more significant carbon stores than was historically thought, given higher-than-predicted productivity and their large surface area, resulting from complex, high-angle topography (Girardin et al., 2014; Spracklen & Righelato, 2014). Thus predictive models of succession may now become essential tools in mitigating the climate crisis: an understanding of seral stages (if indeed they exist!) in forests will be necessary for reaching maximal carbon storage and protecting existing carbon reserves.

The current study was undertaken at Reserva Los Cedros, a mid-elevation cloud forest reserve located in the heart of the Tropical Andean biodiversity hotspot (cite our 2019 paper). While the tropical Andes may not display the same levels of alpha tree diversity often seen in the forests of the lower Amazon basin (Draper et al., 2019; Valencia et al., 1994), the Tropical Andes have long been recognized for a high biodiversity across multiple taxonomic groups (Brehm et al., 2016; Churchill, 2009; Gentry & Dodson, 1987; Hughes, 2016; Küper et al., 2004; Olson, 1994; Ramírez-Barahona et al., 2011, p.; Ríos-Touma et al., 2017; Smith et al., 2007). This biodiversity is characterized by endemism at very fine spatial scales (Borchsenius, 1997; K. R. Young et al., 2002), making it an extremely important conservation priority (Myers et al., 2000). The mechanisms for the high biodiversity and endemism in the tropics-at-large, including montane forests, have been debated for decades and perhaps centuries(Janzen, 1967; Von Humboldt & Bonpland, 2010). High solar energy and constant, plentiful rainfall are characteristic in many areas of the neotropics, and these traits are often associated with high biodiversity (Hawkins et al., 2003). Additionally, the neotropics that may have higher speciation rates, acting as evolutionary “cradles” and/or have higher retention of taxa, thus acting as refugia or “museums”, allowing for greater accumulation of species over geological time (Arita & Vázquez-Domínguez, 2008; Jablonski et al., 2006). The tropical Andes augment these general biodiversity trends in the tropics with characteristics that promote even further the role of endemism: complex topography creates diversified habitats and environmental gradients and increased niches(Gentry, 1992; Hughes, 2016; Janzen, 1967), increasing the potential for sympatric speciation, and also adding dispersal barriers, increasing the potential for fine-scale allopatric speciation (Särkinen et al., 2012). The insular or “island” nature of cloud/mountain systems may be particularly enhanced situations for speciation events that result from neutral events, especially founder effects (Gentry, 1992). All of these factors presumably interact and sum to create the unique patterns of diversity and endemism observed in the Andes.

Here we examine the dynamics of early (<20 years) forest succession at Reserva Los Cedros, following anthropogenic or natural-gap-forming-disturbances. The landscape of the reserve is dominated by primary forest, but contains mixed-age secondary forests of varying land use histories (see results). In regions where remnants of ancient forest ecosystems - ecosystems without a history of significant modern anthropogenic disturbance - persist on the landscape, there are advantages for both ecological analysis and restoration efforts.. First, we have information in the form of local, functioning examples of the biological complexity and reference points for primary-forest equilibrium. Second, forest recovery is facilitated by the presence of species that have co-evolved in local conditions for many millennia, available to act as nuclei for reforestation efforts (Corbin & Holl, 2012; Zahawi et al., 2013). The existence of extensive primary forest has proven particularly valuable for the study of succession and community equilibria in neotropical forests (Bhaskar et al., 2014; Loughlin et al., 2018; Norden et al., 2009).

We also examine the spatial signature of this fine scale beta diversity of trees, with the working hypothesis that individual, steep-sided catchments may be the spatial unit of importance for understanding Andean biodiversity. Our hypothesis was that two characteristics/processes govern the behavior of community similarity in the tropical Andes: (1) large-scale spatial auto-correlation will cause short-distance comparisons to be more similar than farther comparisons, and with with distance, all site comparisons will approach complete dissimilarity, a phenomenon known as Tobler’s Law (Tobler, 1970). However, (2) we predict significant “noise” around this general trend of decreasing similarity, because complex topography of the Andes causes great dissimilarity even among some very localized comparisons, and conversely, causes highly similar habitats to occur sometimes at great distances apart due to similar site conditions. We predict that in systems with extremely complex topography such as the Andes, the most informative unit for modeling community dissimilarity will not be euclidean distance (meters), but instead the number of watersheds crossed. If this hypothesis is supported, it will facilitate future understanding of tropical diversity.

Finally, Los Cedros has succeeded at both forest protection and reforestation, despite its location in a region of high deforestation and now mineral exploration. Thus we examine here the success of los Cedros in conservation of forest in a region of Ecuador that is under intense extractivist pressure, in terms of forest cover change and IUCN red-list species observed.

## Methods

### Site – Reserva los Cedros

All fieldwork described was performed at or directly adjacent to Reserva Los Cedros, a protected forest reserve in the western slope of the Andes, in northwestern Ecuador (00°18 0 31.0 00 N, 78°46 0 44.6 00 W), at 1000-2700 m asl. The reserve lies within the Andean Choco bioregion, one of the most biodiverse habitats on the planet (Gentry, 1992). It is also considered to be within the Tropical Andean Biodiversity hotspot (Myers et al., 2000). The reserve protects 5256 hectares of cloud forest. Definitions of cloud forest vary an can be quite complex (Stadtmüller, 1987), but following Foster (2001) and Stadtmüller (1987), we use the term cloud forest to mean a forest “whose characteristics are tied to the frequent presence of clouds and mist,” and consider the terms montane rain forest and cloud forest to be synonyms. Following Grubb et al. (1963; Grubb & Whitmore, 1966), the ecotones within the reserve are probably most accurately classified as cloud forest of mostly lower montane rain forest, with some regions of higher montane cloud forest, and some Elfin forest zones in its highest, least-explored areas. The Reserve also shares a border with the 305,000 hectare nationally-protected Cotocachi-Cayapas Ecological Reserve. Rainfall averages 2903 (+/- 186 mm) per year according to onsite measurements. Humidity is typically high (∼100%), and daily temperatures at the site range from 15°C to 25°C (Policha, 2014). Annual fluctuations in temperatures are minimal. Daily precipitation varies according to the time of year, with the wet season (October to May) to dry (June-September) seasons (___Ecuador weather service, Nanegalito and Garcia Moreno weather stations, accessed by WeatherAtlas__).

The rarity of primary forest in the north Andean region is due to deforestation from pre-colonial times (Loughlin et al., 2018; Wunder, 2000), and more recently in the 1960’s, due to land reform efforts that legalized and encouraged homestead-scale settlement of large government and private (“Hacienda”) forest holdings (Dodson & Gentry, 1991). Los Cedros has apparently largely escaped deforestation during both eras. The land within Los Cedros and the surrounding region was inhabited by the poorly understood Yumbo indigenous group until 1690.. Their activities probably significantly altered sites but with unknown effects on the ecology and canopy cover of the forest, but their economy was likely integrated into the forested setting of the mid-elevation Andean region (R. D. Lippi, 1988; R. Lippi & Gudiño, 2010). However, the impacts of indigenous land use in South American forests has been until recently been greatly underestimated and mis-understood by scholars (Loughlin et al., 2018; Watling et al., 2017). Given the rainfall and high humidity it is unlikely that large-scale deforestation resulted from fire at Los Cedros. Other than possible small-scale pre-colonial indigenous activities, the majority of the Los Cedros forest has seen relatively few anthropogenic alterations.

Land acquisitions to build the reserve were made between 1988-1995.. Tracts purchased by Los Cedros were usually developed or deforested only in small proportions of their total area, usually for small-scale mixed (“finca”) agriculture or for cattle and mule pasture. In some cases, these small-scale clearings were made for assertion of legal rights over land rather than agricultural production at scale (Jose DeCoux, pers. comm.). Following their acquisition by Los Cedros, all former agricultural and pasture sites were selectively grazed for several months by cattle, to reduce competition from graminoids for protected tree seedlings. Tree seedlings were not planted, but were allowed to reestablish by encroachment from adjacent forest and other sources during and after selective grazing. This method is known informally by some workers in the region as “reforestation by cattle rotation”, and is intended to release naturally regenerated tree seedlings from intensive competition from pasture grasses (Goosem et al., 2013) without the use of herbicides.

Satellite data of forest cover from 1990 shows ∼96% forest cover of los Cedros (see results), of which at least 80% is thought to have been primary forest (Jose DeCoux, pers. comm.). Non-forest land use was concentrated in the southeastern portion of the reserve, where the vegetation surveys were undertaken.

### Tree survey and plant identification

#### Selection of sites and categorization of land use history

Primary forest was defined as those forests which have no historical record and no physical evidence of canopy removal or other large structural alteration by humans. However, forest ecosystems exist continuously as matrices of various states of non-anthropogenic gap formation and closure (Chambers et al., 2013; Espírito-Santo et al., 2014; Franklin et al., 2002; M. O. Hunter et al., 2015; Nicholas Brokaw, 1982), including montane wet forests and cloud forests (Crausbay & Martin, 2016). As such, primary forest is given three categories of land cover based on their gap characteristics:

> **Bosque cerrado** (“BC”) = Mature forest, with no physical sign of gap-forming disturbance. Canopy is closed.

> **Bosque secundario** (“BS”) = Sites with evidence of recent gap, now with a closed canopy,

> **Claros del bosque** (“CLB”) = recent, natural gaps in the forest.

In addition to primary forest, sites with histories of anthropogenic disturbance were placed into two categories:

> **Regeneración de fincas agricultura y ganadería** (“RG”) = abandoned small family farms, with land use mostly consisting of pasture maintained for cattle.

> **Regeneración cañaveral** (“RCA”) = Intensively farmed sites, used for sugar cane production.

#### Survey methods

Site selection - surveys were undertaken at Reserva Los Cedros from the years of 2005 and 2007. The southern area of Los Cedros was divided into 4 areas of study, which were then further divided into three sub-blocks each, from which one sub-block was randomly selected and searched for natural gaps. Once located, each natural gap was also accompanied by a BS and BC site, at a minimum of 40 m distance between survey sites. Two additional smaller blocks were added in the vicinity of previously settled areas, to increase coverage of anthropogenic disturbances. Overall 61 sites from various land use histories and elevations were sampled (see fig. 6).

Adult trees were defined as trees at the height 1.5m with a diameter of 10 cm or greater. Material from all trees fitting this description were sampled, within a circle plot of 30 m^2^ radius from the centerpoint for morphological identification to species, where possible. Additionally, within each survey site of BC, RCA, and RG plots, all juvenile trees were examined. Juvenile trees were defined as trees with a maximum diameter less than 10 cm, growing from 50 cm up to 2 m, of species known to be capable of growing to adult tree size as given above, under ideal circumstances and with sufficient time. Juvenile trees were surveyed within a square subplot of size 5m x 5m, centered within the larger 30m circular plot. Nomenclature of identifications is based on the Flora of Ecuador (Persson et al., n.d.).

### Statistical and informatic methods

All analyses were conducted in Python and R. Python version 3.8.10 (Python Core Team, 2018), using Pandas 1.1.3 (McKinney, 2010; Reback et al., 2020) and Matplotlib 3.1.2 (J. D. Hunter, 2007) for visualization. R version 3.6.3 (R Core Team, 2020) with in-box R plotter engine was used. All analysis was in an Ubuntu 20.04.2 LTS environment. Where Bayesian analyses were used, Python version 3.7.3 was used for compatibility with the PyMC3 package, version 3.8 (Salvatier et al., 2016). All Bayesian models used the default NUTS sampler to sample the posterior. Code for all pertinent statistical analyses have been run and recorded in a Jupyter Notebook that is stored in the affiliated [github repo](https://github.com/danchurch/losCedrosTrees/tree/master/anaData), and is viewable as a notebook online [here](https://nbviewer.org/github/danchurch/losCedrosTrees/blob/master/anaData/MariscalDataExploration.ipynb).

#### Species accumulation curves and richness estimators

Species accumulation curves for the entire area studied were calculated as the number of adult tree species observed per meter-squared of physically sampled area. Each subplot covered a circle with a diameter of 30m, or 0.071 ha. Additionally, a permanent tree diversity plot was established concurrently with rapid surveys in 2005. Within this, trees were sampled in a grid format, every 10 m along a east/west axis and every 5 m along a north/south axis, in a rectangular half-hectare area (50m x 100m). Trees were identified to species where possible, and unidentified trees were grouped into species-level operational taxonomic designations (supp. table 1A). In the case of the permanent plot, tree species accumulation was modeled as a function of trees-examined. Species accumulation models and species estimators were calculated using the specaccum, specpool and associated functions with the Vegan package in R, version 2.5-6 (Jari et al., 2019). Point predictions of diversity for 1 ha area were generated using the predict function, which when used as a method of specpool model objects uses a Mao Tau rarefaction method (Colwell et al., 2012) to generate predictions.

#### Tree community turnover (distance decay or beta-diversity)

Turnover in tree communities was modeled and visualized using Bray-Curtis dissimilarity (Bray & Curtis, 1957; Legendre & Legendre, 2012) as a function of (1) physical distance or (2) watershed crossings mapped at small-scale. Bray-Curtis dissimilarities and physical distances matrices among subplots were calculated using the SciPy spatial submodule (Virtanen et al., 2020). Bayesian models of community dissimilarity decay were conducted using PyMC3 package in Python, with visualizations created using the companion ArViz package (Kumar et al., 2019).

##### Turnover by physical distance

Tree community turnover by physical distance was calculated using Bray-Curtis dissimilarity as a function of physical distance (meters). Two Bayesian models were used as alternative formulations of the above two processes of local variability and large-scale spatial autocorrelation.

1. **Asymptotic model**: In one approach, BC dissimilarity was modeled as an asymptotic function, or “Michaelis-Menten”-type function:

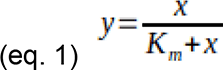

Where y is predicted Bray-Curtis dissimilarity, x is distance between compared sites, and K_m_ is the distance at which half of maximum BC dissimilarity is reached. This model honors both theoretical constraints of complete similarity at proximal sites and maximum dissimilarity. Additionally, we modeled variation around the mean as a linear variable, reducing as all distant comparisons approached complete dissimilarity (BC=1). This shrinking variance term ɛ was exponentiated to avoid negative values. As a Bayesian model this is formulated in probabilistic terms, with priors, as follows:

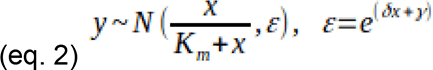

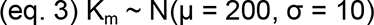

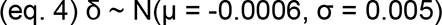

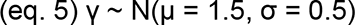

Prior distribution for K_m_ was loosely estimated from Draper et al. (2019), who showed that in many Amazonian forest ecosystem types, decay of tree community similarity occurs rapidly, losing more than half their similarity within 200 m. Owing to the novel formulation, priors for delta and gamma of equation 2 were not found in existing literature, and were assigned weak priors, with means intended to reflect the initial wide variation and its subsequent tightening around BC=1.

2. **Linear model**. In a second formulation, decay in tree community similarity was modeled as a simple linear equation. Variance around the mean BC dissimilarity was still allowed to vary negatively with distance, beginning with large initial spread to encompass both highly similar and dissimilar sites at close distances. Additionally, a skewed mean distribution (Ashour & Abdel-hameed, 2010; O’Hagan & Leonard, 1976) was used to describe the variance around the mean function, given the long skew that is observable in the data at most distances. As with the asymptotic model, the variance around the mean function of BC dissimilarity was described as an exponentiated linear function.

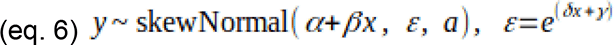

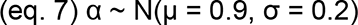

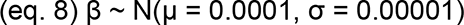

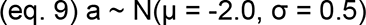

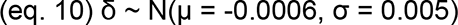

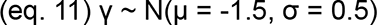

Prior distribution for β was again loosely estimated from Draper et al. (2019), using data from community turnover of Amazonian Terra firme forests. All other priors were assigned as weakly informative priors.

###### Model comparison

Comparison of performance between these two models were done using posterior predictive checks on the data and variance explained (Bayesian R^2^), using the sample_posterior_predictive command in the PyMC3 package, and the r2_score command in the ArViz package. Bayesian p-values to indicate balanced performance were calculated using the distribution of residual differences between model predictions and observed data. To confirm Bayesian linear model results, a classic linear least-squares regression was also applied to the data, using a Wald test with a null hypothesis that the slope of community turnover was zero, as implemented with the linregress function in the SciPy Stats module. Variance defaults (bivariate normal) for the function were not modified.

##### Overall turnover by watershed crossings

To better understand changes in community as a function of topographic complexity, we delineated small watersheds in the area of the study area. The original digital elevation model used was the ASTER Global Digital Elevation Map version 2 (Tachikawa et al., 2011). Every subplot was assigned to a micro-watershed, and a distance matrix of watershed crossings. Watersheds were delineated using the pysheds package in Python (Bartos, 2021). Distance matrix of micro-watershed crossings was calculated using Djikstra’s algorithm for shortest paths (Dijkstra, 1959) in the NetworkX package, version 2.4 (Hagberg et al., 2008). Methods for specific watershed delineations are further explained in the statistical jupyter notebook (https://nbviewer.jupyter.org/github/danchurch/losCedrosTrees/blob/master/anaData/MariscalDataExploration.ipynb#watersheds).

#### Ordination by historical land-use/habitat

Structuring in adult tree communities as a function of historical land-use/habitat by was visualized using Bray-Curtis dissimilarity and non-metric multidimensional scaling (NMDS) (Legendre & Legendre, 2012), using the metaMDS function in the Vegan package in R.

#### GIS data and additional environmental data

Except where otherwise noted, geospatial data were explored and visualized using using tools from the GeoPandas (version 0.8.1, https://geopandas.org) and Rasterio (version 1.1.8, https://rasterio.readthedocs.io/), and QGIS (QGIS v3.4.11-Madeira, OSGEO v3.7.1). Several environmental variables were generated from ASTER Global Digital Elevation Map version 2, including slope, aspect, elevation, eastern and northern exposure, and distance-to-nearest-stream of all subplots. Stream data were digitized from Map CT-NII-C3-d of Bosque Protector Los Cedros, from the Ecuadorian Instituto Geográfica Militar. Rasters of slope and aspect were generated with the Raster Terrain Analysis tool suite in QGIS.

#### Hierarchical clustering of sites

To understand the current types of forests present at los Cedros, adult tree community data from all sites were partitioned into clusters using Ward’s minimum variance clustering (Legendre & Legendre, 2012) using Bray-Curtis dissimilarity, as implemented in the hclust command in the Vegan package in R. Number of clusters was decided by eye from the resulting dendrogram, taking into consideration branch lengths and meaningful ecological groups.

#### Prediction of current ecological state by land use history and elevation

Current ecological state as defined from cluster analysis was modeled as a function of historical land-use/habitat data and elevation. Each of the two predictors (height, land-use/habitat) was considered individually, and then a combined model was created using both elevation and historical land-use/habitat. All models were Bayesian, written in python with the PyMC3 package.

**For land-use/habitat**, a multinomial logistic regression model was constructed with a softmax link function and the following priors:

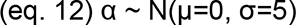

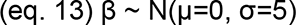

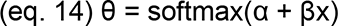

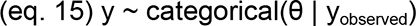

Where y is a vector of predicted probabilities for forest type (cluster) of a site, distributed as a categorical random variable of α, β, and x, conditioned on our observed current forest type (i.e., cluster number). x is observed historical land-use/habitat, in a dummy variable matrix format. α are y-intercepts for each current forest type, and β is the vector of slope coefficients for each of 5 types of historical land-use/habitat, for each current forest type. α and β were assigned weak, normally distributed priors centered on zero.

##### Elevation

A logistic regression model was constructed to model differences among forest types III and IV from the hierarchical cluster analysis. Hierarchical clustering of tree communities and NMS ordinations both indicated two distinct “natural” forest types occurring within the extent of the survey, which were designated Forest types III and IV (see results). To further understand differences between these two forest types, exploratory PERMANOVA models were used to test grouping by all available environmental predictors (see Jupyter notebook). Following this, forest type was modeled as a function of elevation, using a Bayesian logistic regression model. Priors and posterior were modeled as follows:

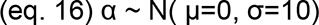

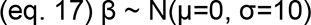

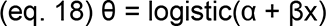

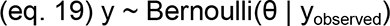

Where α and β are the intercept and slope controlling the boundary decision between type III and type IV forests, in terms of elevation. θ is the prior probability that a forest site will be type III forest, given its elevation, and y is a Bernoulli distribution with θ as the mean, conditioned by the observed forest type.

##### For the combined elevation and historical land-use/habitat model

Elevation and historical land-use/habitat predictors were combined in one linear model:

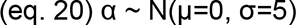

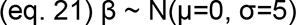

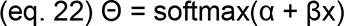

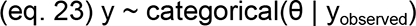

Where x is observed historical land-use/habitat, in a dummy variable matrix format, with an additional column vector giving the elevation of each site. α are y-intercepts are intercepts for each row of x, and β is the vector of slope coefficients for each column of x for each current forest type (= row of x). α and β were assigned weak, normally distributed priors centered on zero. y is a vector of predicted probabilities for forest type (cluster) of a site, distributed as a categorical random variable of α, β, and x, conditioned on our observed current forest type (i.e., cluster number).

#### Indicator species analysis

Indicator species analyses were conducted to ascertain representative species for both historical land-use/habitat types and current forest type (cluster analysis results), using the indicspecies package in R (De Cáceres et al., 2010), using the multipatt function and Pearson’s phi coefficient correlation (function “r.g”).

#### Spatial analyses

To determine important spatial scales on which tree community was changing, and to give shape to possible spatial structuring of tree community, Moran’s eigenvector maps (MEM) were constructed (Dray et al., 2012), following examples as found in documentation for the ADEspatial package (https://cran.r-project.org/web/packages/adespatial/vignettes/tutorial.html). A spatially-weighted neighborhood matrix of sample sites was constructed using a Gabriel connectivity matrix with an inverse distance weighting. Abundances of our adult tree community matrix were Hellinger transformed (Legendre & Legendre, 2012), and new PCA axes of the transformed were determined with the dudi.pca command in ade4 package. Important MEMs were then selected on the basis of their p-value as explanatory terms in a linear model with the 4 most important tree community PCA axes as response variables. Important MEMS were determined using a forward model selection process, as implemented with the mem.select command in the adespatial package. The default cutoff of alpha = 0.05 was used to decide statistical significance of MEMs. All available environmental variables were checked for correlations with MEMs using standard linear regression as implemented in the linregress function in the Python SciPy Stats package. Correction for multiple testing was done with a Benjamini-Hochberg correction, as implemented in the multipletests command in the Python Statsmodels package. For exploratory purposes, the false discovery rate was set at FDR=0.1.

#### Juvenile communities

Juvenile tree community data were available for a subset of historical land-use/habitat types: “Bosque Cerrado” (CB), “Regeneración cañaveral” (RCA), and “Regeneración fincas agricultura y ganadería” (RG). Juvenile data were not taken for “Claro del Bosque’’ (CLB) or “Bosque secondario (“BS”). Due to the difficulties associated with identifying poorly understood herbaceous taxa and/or juvenile trees, these data required extensive additional cleaning and verification, following botanical identifications. For the purposes of this study, juvenile trees were defined as undersized plants that would, in time and with proper growth conditions, reach the stature of the mature trees as defined above, namely: woody plants with a diameter-at-1.5m of 10 cm or greater. This is as compared to herbaceous plants or small woody plants (such as lianas or subshrubs) which would never or rarely reach 10 cm girth at 1.5m height.

Samples that did not receive sufficient identification to confidently conclude that the sample was indeed a juvenile tree were discarded from the analysis. This selection was done by automated and manual queries of growth form data requested from the TRY database (Kattge et al., 2020), manual checks of the Encyclopedia of life (Parr et al., 2014), and the Smithsonian Tropical Research collections site (https://stricollections.org/portal/index.php). The Gentry manual (Gentry, 1993) was also frequently consulted. Exact species information was not often available and designation of juvenile tree status was made using genus growth form data where possible. Juvenile tree communities were compared to adult trees by subsetting adult tree community data to just those sites with juvenile data, and combining community matrices from both. These were then ordinated using Bray-Curtis dissimilarity and non-metric multidimensional scaling (NMS), using the metaMDS function in the Vegan package in R.

#### Deforestation in the region

To understand the extractive pressure in the region of Los Cedros, and the efficacy of Los Cedros as a conservation project, we examined land cover changes from 1990 to 2018. Data for changing land cover were accessed from the Ecuadorian Ministerio del Ambiente’s (“MAE”) interactive map and GIS catalogue, using the “Cobertura vegetal” layers from 1990 and 2018, available [here](http://ide.ambiente.gob.ec/mapainteractivo/). Vector data from the MAE were rasterized to a pixel resolution of 30 m^2^ using the gdal_rasterize program from the GDAL tool suite from the OSGeo project (GDAL/OGR contributors, 2021), and visualized in Matplotlib using the rasterio package. Background deforestation rates were constrained to Cotacachi canton of the Imbabura province in which Los Cedros is located. Deforestation rates in the similarly-sized, nearby Bosque Protector el Chontal were quantified for comparison with los Cedros. BP Chontal is managed by the Cattlemen’s Association of the nearby town of Brillasol (pers comm. Jose DeCoux). Administrative boundaries of BP Los Cedros and BP Chontal were supplied using official boundary layer “Bosque y Vegetación protectora”, also available at MAE’s [interactive map website](http://ide.ambiente.gob.ec/mapainteractivo/).

## Results

Particularly interesting or pertinent posterior distributions are described here. Full results on posterior distributions of all models are available in the [jupyter notebook](https://nbviewer.jupyter.org/github/danchurch/losCedrosTrees/blob/master/anaData/MariscalDataExploration.ipynb).

### Species accumulation curves and richness estimators

343 adult tree species were directly observed in plots. Estimates of total adult tree species in the study area range from 404 to 566 species (fig. 1A, table 1). In the permanent 0.5 ha plot, 43 species of tree were observed, distributed among 36 genera and 25 plant families, with a range of predicted total species from 51 to 72 species in the half-hectare (fig. 1B, table 1). On average, 1 ha in the southern portion of Los Cedros is predicted to host 169 species of adult tree. Chao estimates, first and second order Jackknife, and Bootstrap estimates for the total study area and for the permanent 0.5 ha plot are given in table 1.

**Figure 1, A,B.**
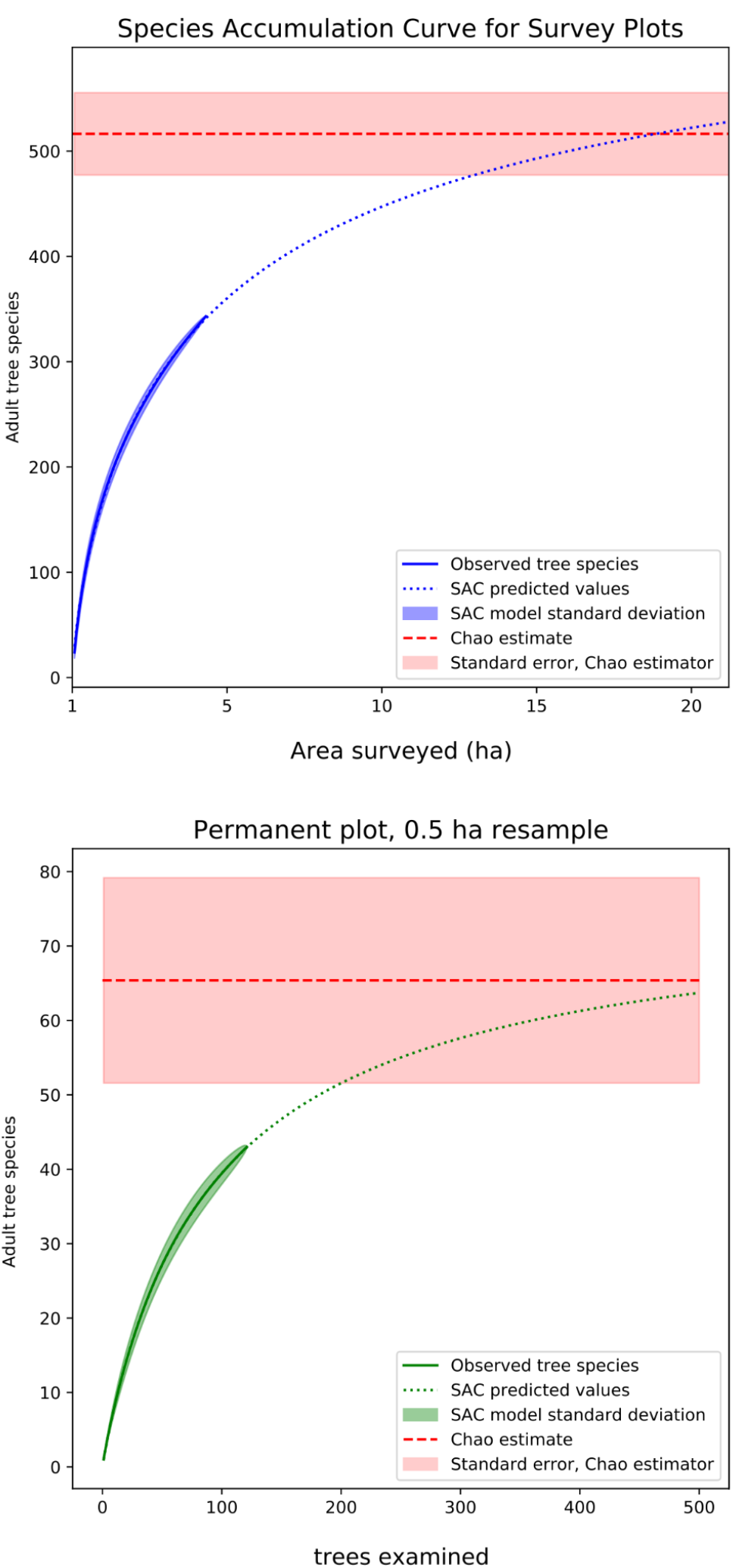

**Table 1.** Estimates of adult tree species diversity in survey area and permanent plot.

|  | sample size | species observed | chao | chao.se | jack1 | jack1.se | jack2 | boot | boot.se |
| --- | --- | --- | --- | --- | --- | --- | --- | --- | --- |
| Survey Plots | 61 | 343 | 516.4 | 39.2 | 483.7 | 21.6 | 566.8 | 404.4 | 10.5 |
| Permanent Plot (Trees) | 121 | 43 | 65.4 | 13.8 | 61.8 | 4.3 | 72.7 | 51.4 | 2.2 |

When examined by land-use/habitat type, more adult tree species were observed from reforested pasture sites, possibly due to deeper sampling, but natural-disturbance sites (BS, BC, and CLB) were comparable when rarified to a common sample size (fig. 1C, supp. table 2). Sites with history of intensive agricultural use were the least diverse, though depth of sampling was lowest in this group. Species estimators by land-use history are given in supp. table 2.

**Figure 1,C.**
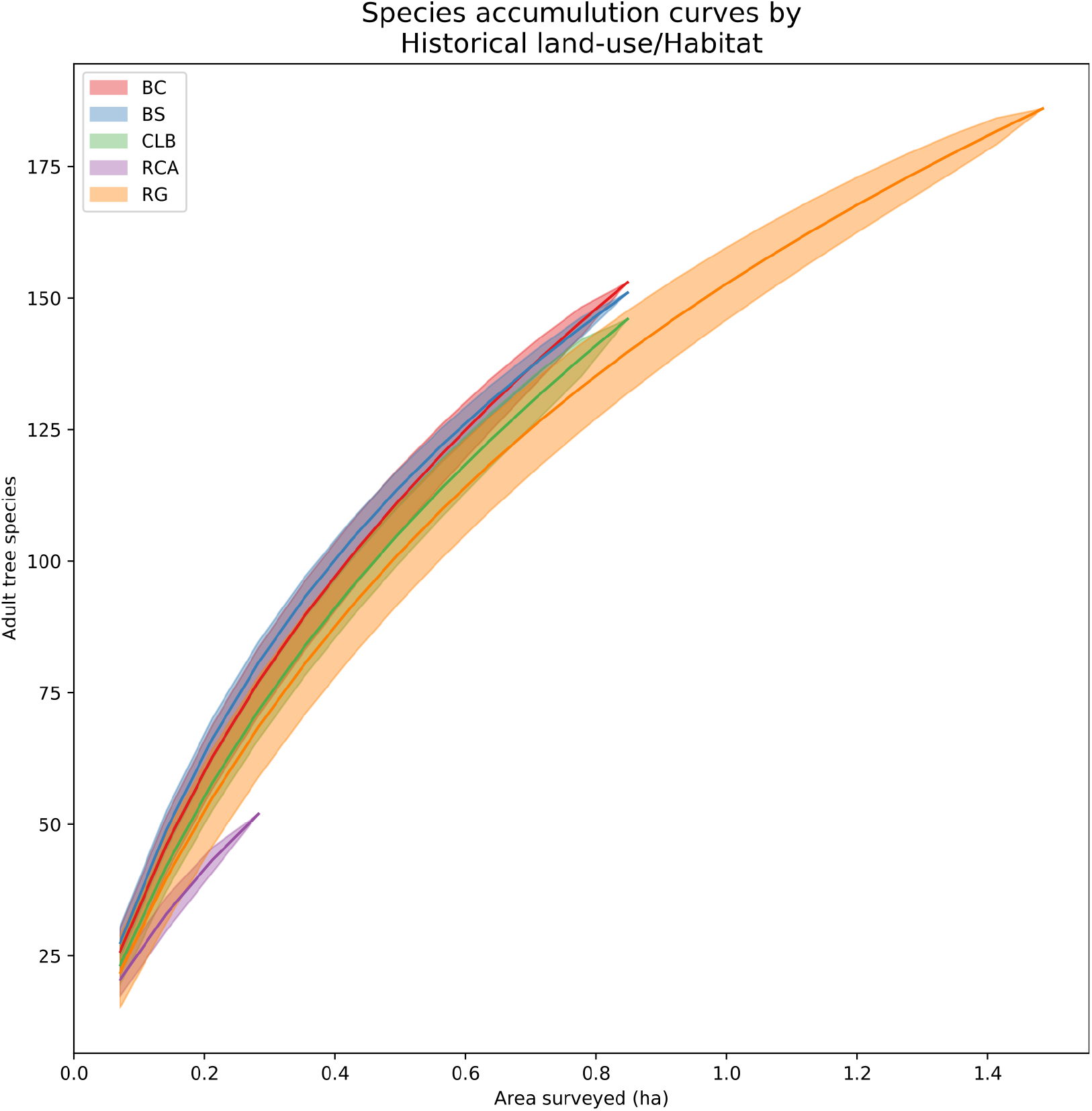

### Tree community turnover (distance decay)

#### Turnover by physical distance

##### 1. Asymptotic model

The asymptote model is visualized with 50% and 95% highest posterior densities (“HPD”) for predicted BC values in fig. 2A. The posterior distribution of our asymptotic model explained a large amount of the variance in the adult tree community data (R^2^ = 0.53 +/- 0.11). The distribution of posterior predictive values was nearly symmetric around observed Bray-Curtis values (Bayesian p-value = 0.49, supp. fig. 1). The posterior distribution of K_m_ was centered on 153 m (+/- 6.13 m), a shift of −47m from the prior estimate of 200m, with no overlap between posterior and prior (supp. fig. 2).

**Figure 2A.**
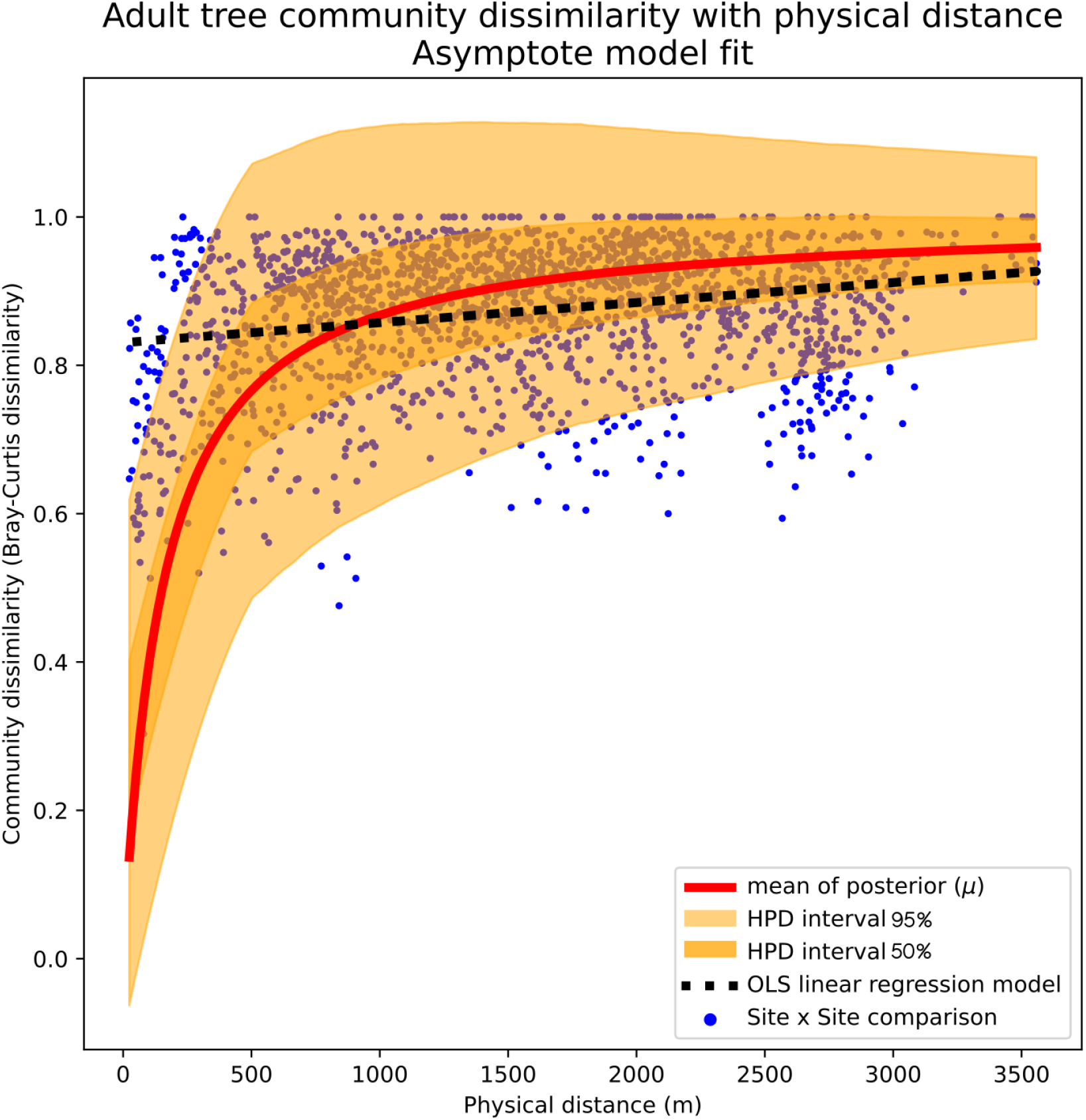

##### 2. Linear model

The Bayesian linear model is visualized with 50% and 95% highest posterior densities (“HPD”) for predicted BC values in fig. 2B. As constructed, the posterior distribution of our linear model explained only a small amount of the variance in the adult tree community data (R^2^ = 0.06 +/- 0.01). The posterior predictive values tended to be lower than observed BC values (Bayesian p-value = 0.41, supp. fig. 1). The frequentist, least-squares linear regression model highly resembled the Bayesian model, also reporting very little variance explained (R^2^ = 0.06 +/- 0.01, p = <0.01, fig. 2B).

**Figure 2B.**
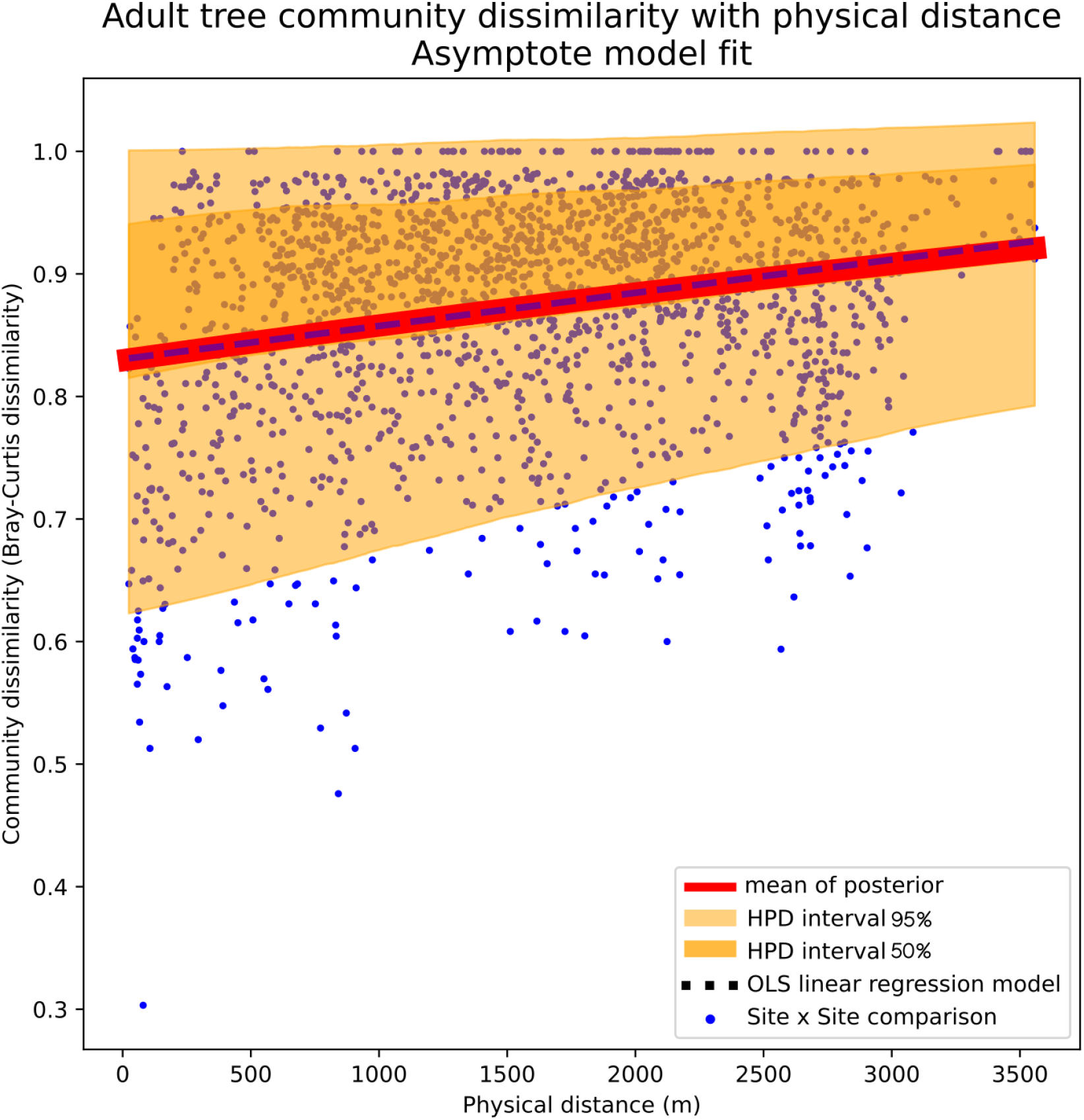

#### Overall turnover by watershed crossings

When plotted as a function of watershed crossings (fig. 3), the widest range of Bray-curtis values occurred among sites within the same watershed (distance class 0), as did the lowest mean BC score (mean BC dissimilarity = 0.82, +/- 0.11). Following this, other mean BC values of comparisons of other distance class did not vary heavily, and are mostly statistically indistinguishable (Kruskal-Wallis H-test (H(5, 1825)=73.27, p<0.01, and Tukey HSD (p.adj=0.001), table 2 and table 3, fig. 3). After the first watershed crossing, further comparisons are all approximately equally dissimilar, with a mean of BC = 0.88 (+/- 0.086).

**Figure 3.**
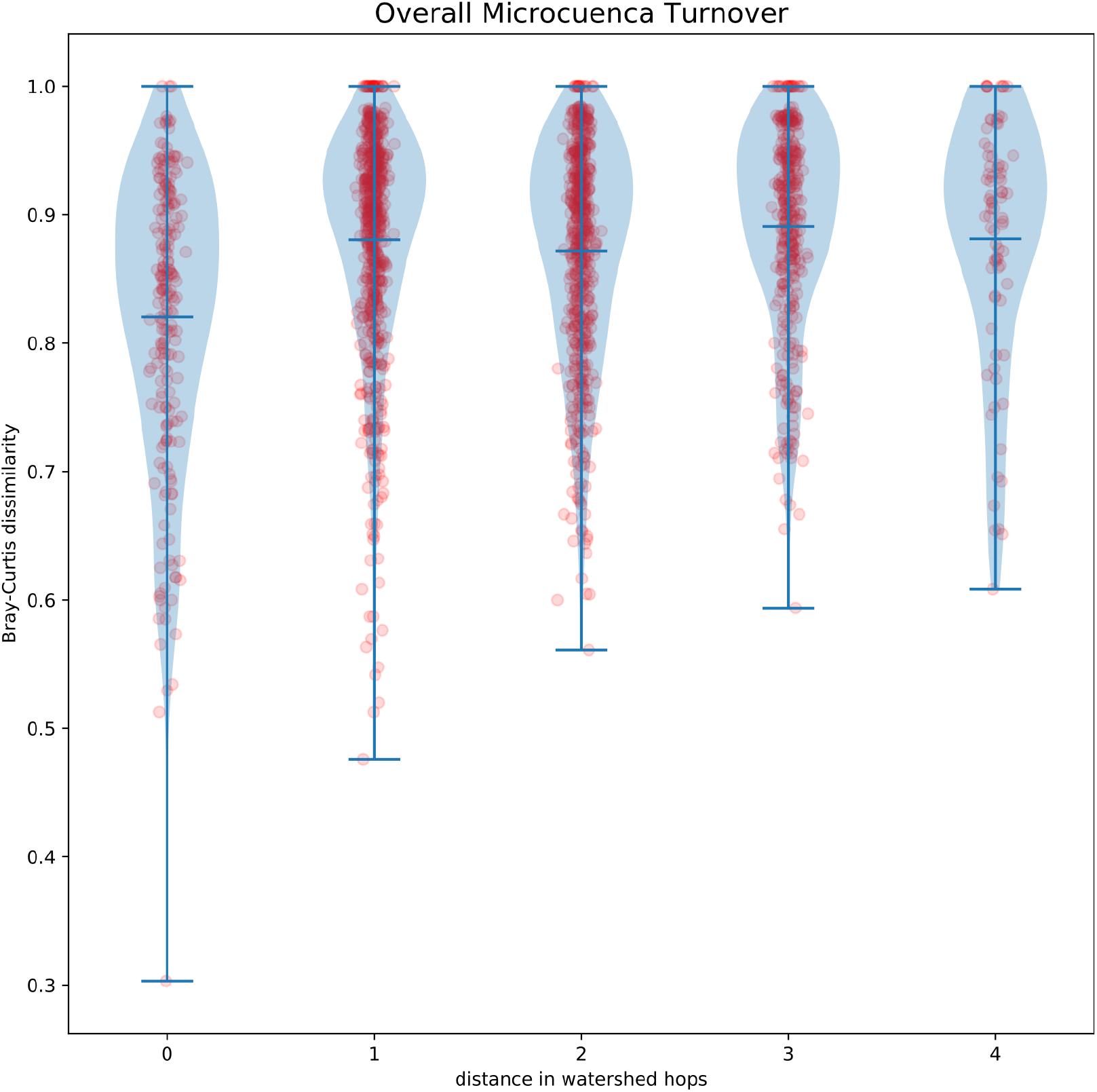

**Table 2.** Mean dissimilarity of survey sites, by number of watersheds crossed.

| watersheds crossed | sample size | mean BC | stdDev BC |
| --- | --- | --- | --- |
| 0 | 215 | 0.820 | 0.114 |
| 1 | 630 | 0.880 | 0.089 |
| 2 | 559 | 0.872 | 0.085 |
| 3 | 346 | 0.891 | 0.079 |
| 4 | 80 | 0.881 | 0.096 |

**Table 3.**
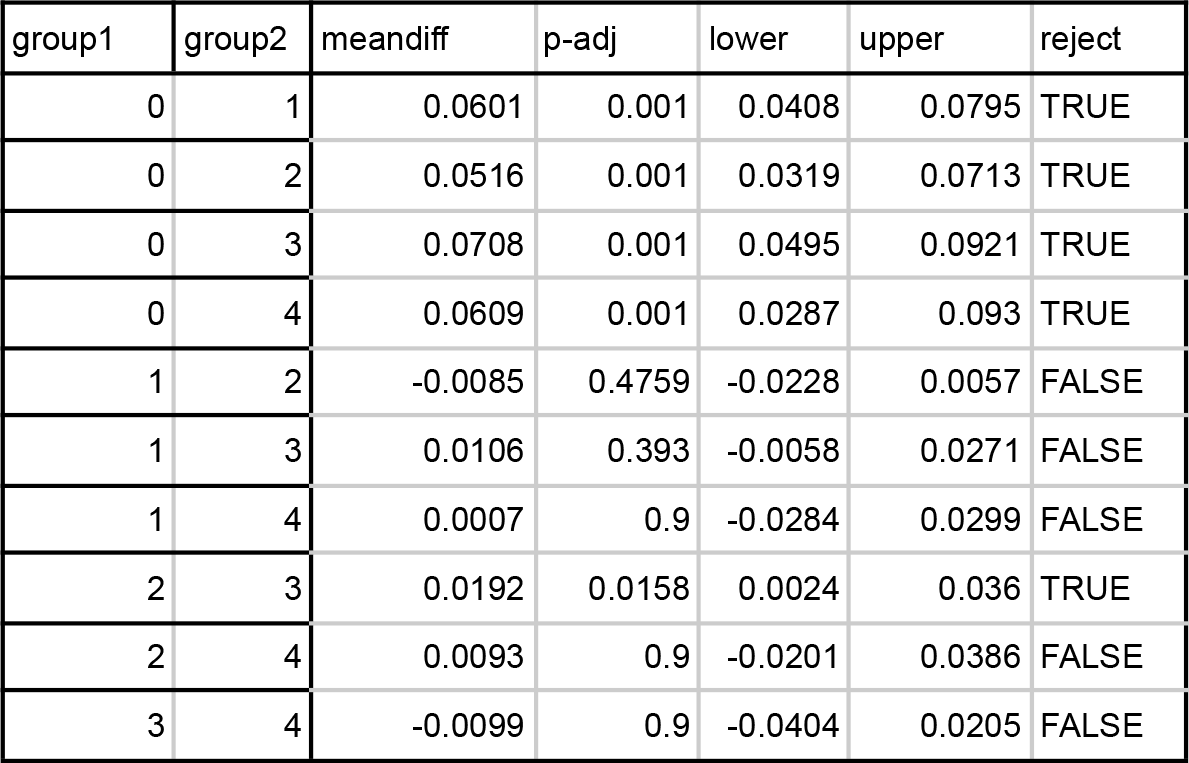
Tukey’s Honest Significant difference of comparisons among watershed distance classes.

| group1 | group2 | meandiff | p-adj | lower | upper | reject |
| --- | --- | --- | --- | --- | --- | --- |
| 0 | 1 | 0.0601 | 0.001 | 0.0408 | 0.0795 | TRUE |
| 0 | 2 | 0.0516 | 0.001 | 0.0319 | 0.0713 | TRUE |
| 0 | 3 | 0.0708 | 0.001 | 0.0495 | 0.0921 | TRUE |
| 0 | 4 | 0.0609 | 0.001 | 0.0287 | 0.093 | TRUE |
| 1 | 2 | -0.0085 | 0.4759 | -0.0228 | 0.0057 | FALSE |
| 1 | 3 | 0.0106 | 0.393 | -0.0058 | 0.0271 | FALSE |
| 1 | 4 | 0.0007 | 0.9 | -0.0284 | 0.0299 | FALSE |
| 2 | 3 | 0.0192 | 0.0158 | 0.0024 | 0.036 | TRUE |
| 2 | 4 | 0.0093 | 0.9 | -0.0201 | 0.0386 | FALSE |
| 3 | 4 | -0.0099 | 0.9 | -0.0404 | 0.0205 | FALSE |

### Hierarchical clustering of sites and prediction of current ecological state by land use history and elevation

Tree communities first sort into two large groups that align well with previous land use - the first grouping contains sites that have historically experienced anthropogenic disturbances of any type, and the second grouping contains sites with no recorded history of anthropogenic disturbance, but all types of natural gap-forming disturbances (fig. 4). Each of these groups then sorts into two further groups (for a total of four clusters). Cluster I contains all sites considered to be highly disturbed by agricultural use (RCA sites), and some sites with intermediate agricultural use as pasture (RG sites) (fig. 4). Cluster II consists entirely of RG sites, and also contains the majority of RG sites. Sites with no history of anthropogenic disturbances fall into two clusters, type III and type IV (fig. 4). Hierarchical clustering results are congruent with NMS ordinations (fig. 5, supp. fig. 3). A summary map giving both current ecological state and historical land-use/habitat is given in fig. 6.

**Figure 4.**
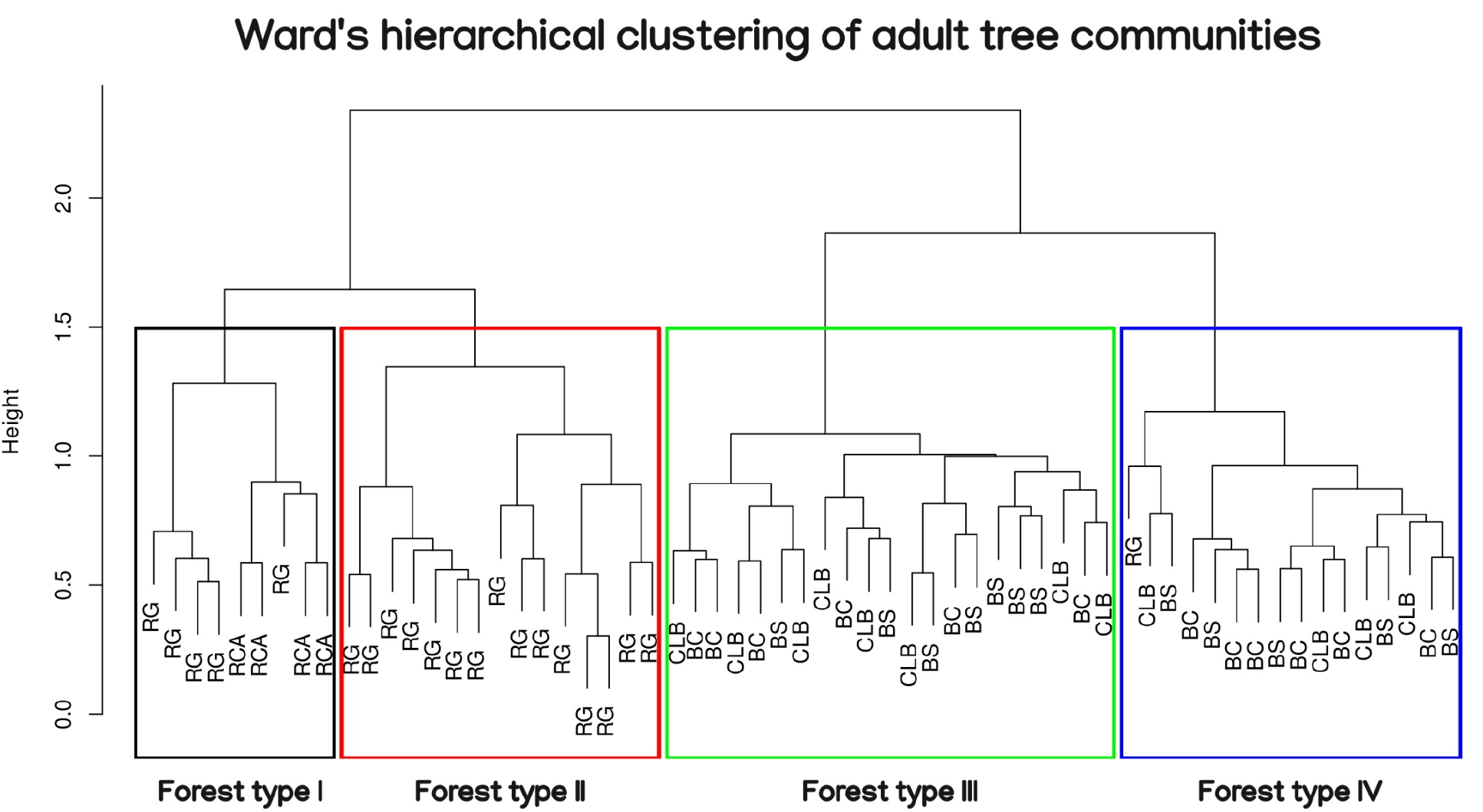

**Figure 5.**
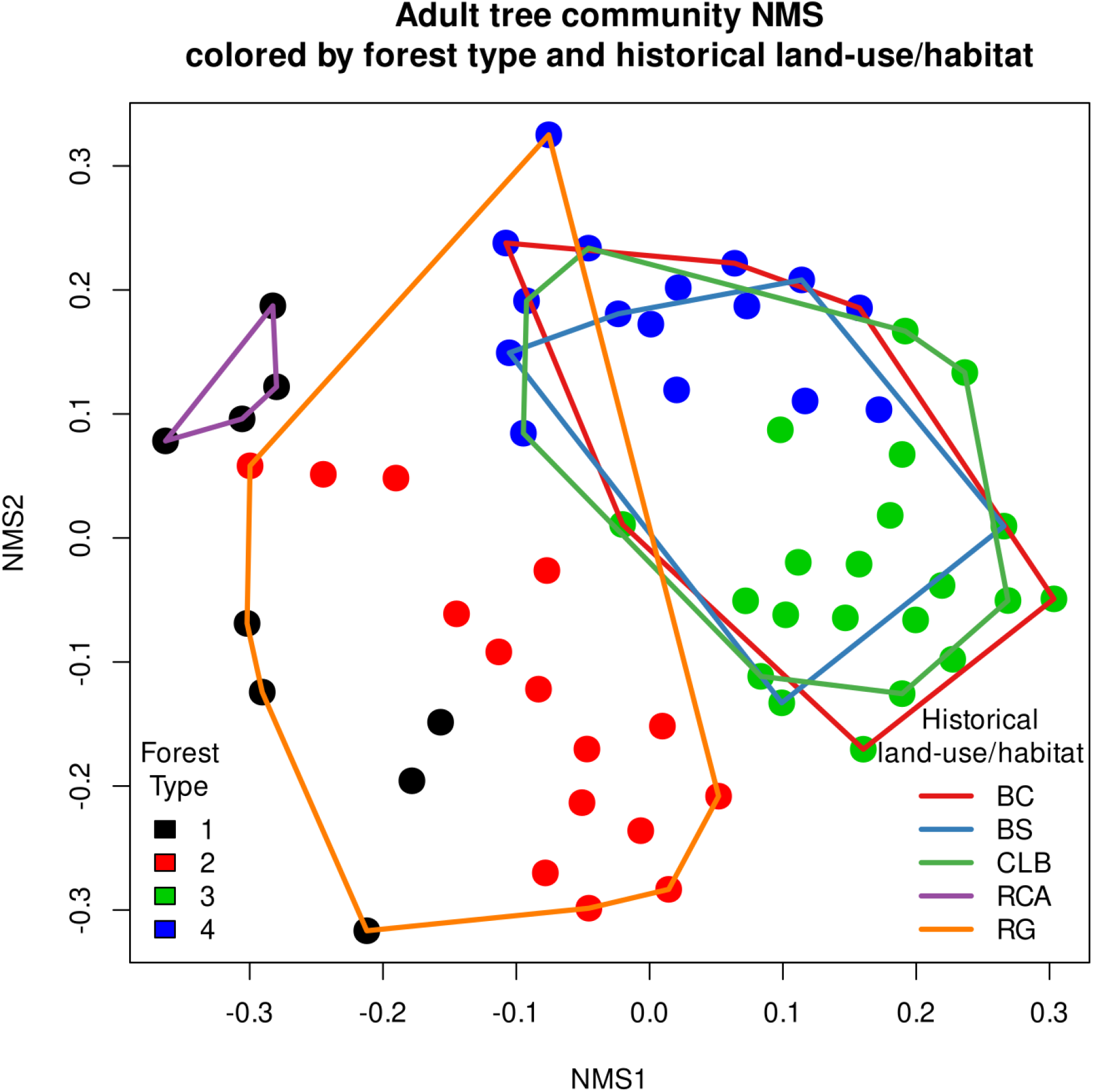

**Figure 6.**
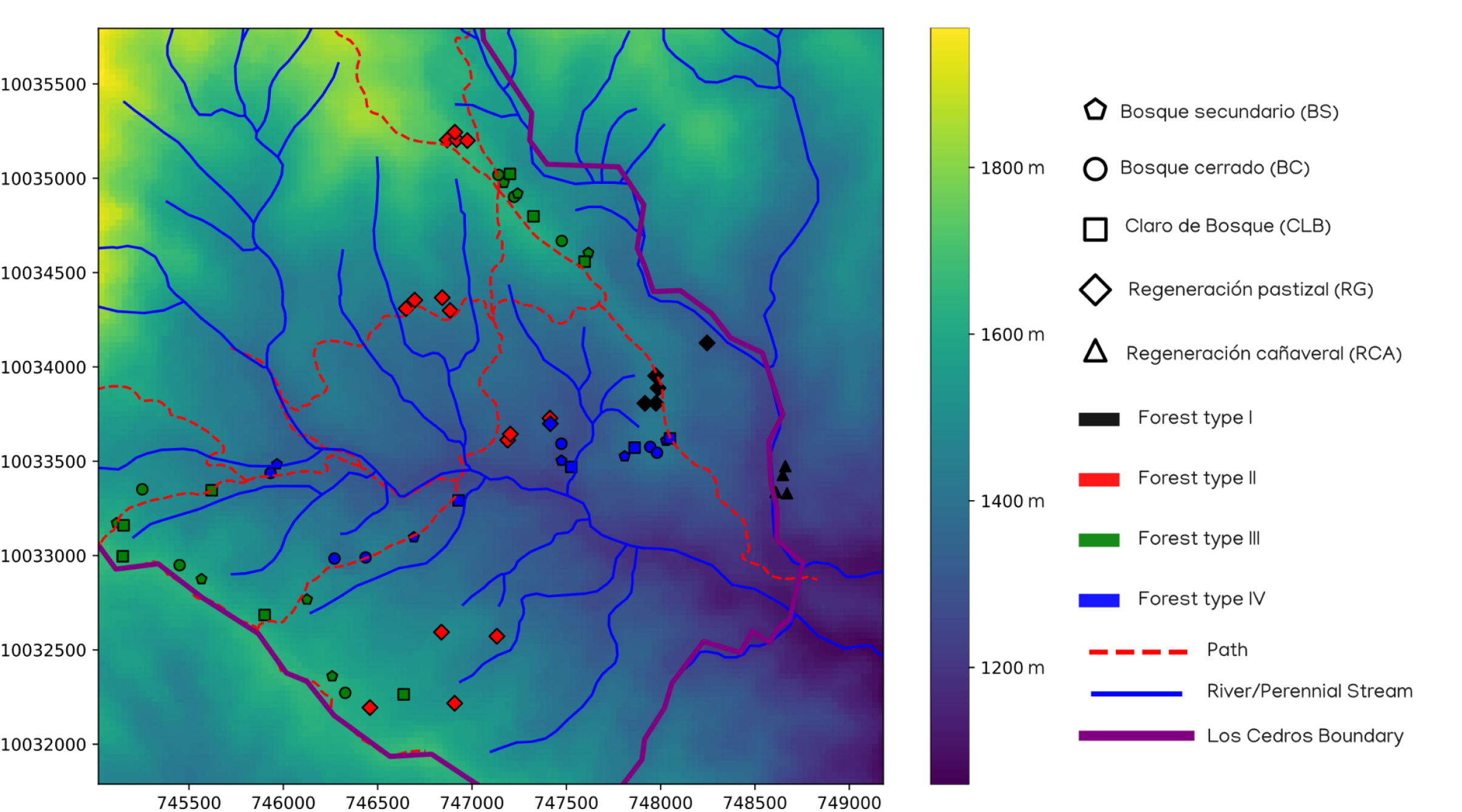

A linear model with these ecological states (forest types I-IV) as response variable, using the single variable of previous land-use/habitat as predictors, successfully predicts the sites that have been modified for pasture or agriculture, and it also well predicts non-anthropogenically sites to types III/IV ecological state (R^2^ = 0.65 +/-0.04, supp. fig. 4). Posterior predictive checks were correct for 65% of sites, mostly for sites with type I and type II forests. However, this previous land-use/habitat-only model cannot distinguish among “natural forest” (III and IV) types.

Among sites with no history of anthropogenic disturbance (“natural forests”, types III and IV), elevation strongly predicts the current ecological category of forest: sites with no history of anthropogenic disturbance at elevations of 1503 m (95% credible interval = 1469 m to 1537 m) or higher were categorized almost completely into type III forests, and below this elevation sites were categorized into type IV forests (bayesian logistic regression, R^2^ = 0.87), fig. 7, fig. 6, supp. fig. 5).

**Figure 7.**
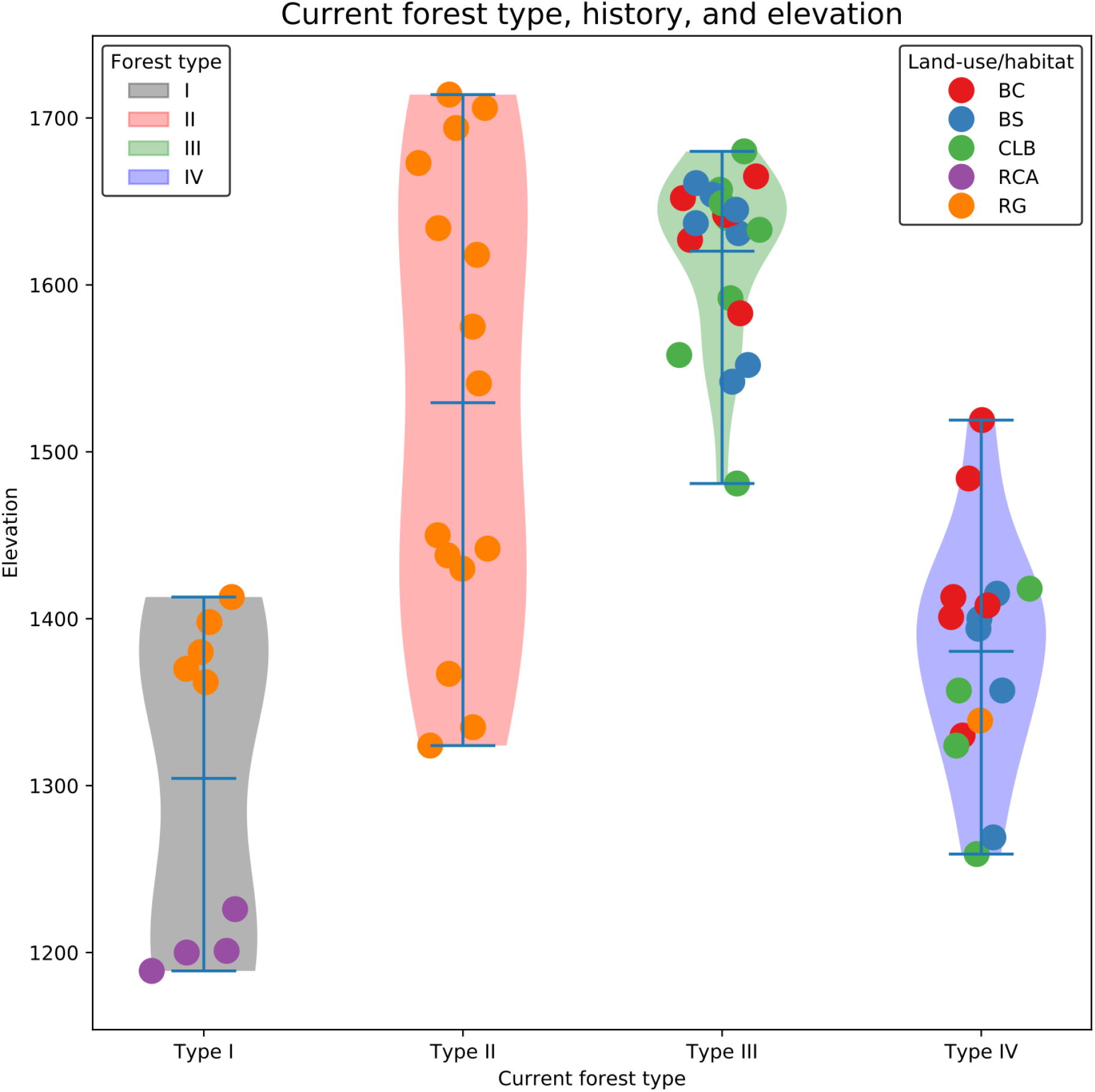

A combined model of previous land-use/habitat and elevation explains most of the variance in total and predicts current ecological state very well (bayesian linear model, R^2^ = 0.88 +/-0.16, supp. fig. 6), and performs much better on posterior predictions, with 87% of sites correctly predicted to their forest type.

### Indicator species analysis

Indicator species were detected for all individual land-use-history/habitat types, all current ecological states (forest types), and for some combinations of site types. Indicator species analysis results are given in table 4A and table 4B.

**Table 4A.** Indicator species for historical land-use/habitat

| Group BC |  |  |  |
| --- | --- | --- | --- |
| species | rpb.g | p.value | sig code |
| Moraceae.Pseudolmedia.rigida | 0.423 | 0.0317 | * |
| Moraceae.Ficus.subandina | 0.406 | 0.049 | * |
| Group BS |  |  |  |
| species | rpb.g | p.value | sig code |
| Rubiaceae.Psychotria.paeonia | 0.478 | 0.0148 | * |
| Lauraceae.Persea.pseudofasciculata | 0.432 | 0.0351 | * |
| Myrtaceae.Myrcia.aff..aliena | 0.425 | 0.0377 | * |
| Group CLB |  |  |  |
| species | rpb.g | p.value | sig code |
| Lauraceae.Endlicheria.cf..chalsea | 0.431 | 0.0268 | * |
| Group RCA |  |  |  |
| species | rpb.g | p.value | sig code |
| Boraginaceae.Cordia.colombiana | 0.894 | 0.0001 | *** |
| Actinidaceae.Saurauia.sp.1 | 0.694 | 0.0003 | *** |
| Melastomataceae.Miconia.aff..brevithec | 0.683 | 0.0004 | *** |
| Moraceae.Ficus.caldasiana | 0.667 | 0.0033 | ** |
| Staphyleaceae.Turpinia.occidentalis | 0.663 | 0.0006 | *** |
| Moraceae.Clarisia.biflora | 0.645 | 0.0022 | ** |
| Lauraceae.Nectandra.membranacea | 0.61 | 0.0027 | ** |
| Verbenaceae.Aegiphila.alba | 0.594 | 0.002 | ** |
| Moraceae.Ficus.andicola | 0.539 | 0.0191 | * |
| Lauraceae.Caryodaphnopsis.theobromifolia | 0.535 | 0.0099 | ** |
| Lauraceae.Nectandra.aff..membranacea | 0.513 | 0.0106 | * |
| Group RG |  |  |  |
| species | rpb.g | p.value | sig code |
| Cecropiaceae.Cecropia.andina | 0.662 | 0.0018 | ** |
| Cecropiaceae.Cecropia.ficifolia | 0.58 | 0.0035 | ** |
| Cecropiaceae.Cecropia.sp.2 | 0.427 | 0.031 | * |
| Melastomataceae.Meriania.tomentosa | 0.415 | 0.0426 | * |
| Group BC+BS |  |  |  |
| species | rpb.g | p.value | sig code |
| Lauraceae.Aniba.aff..hostmanniana | 0.411 | 0.0471 | * |
| Group RCA+RG |  |  |  |
| species | rpb.g | p.value | sig code |
| Solanaceae.Solanum.lepidotum | 0.529 | 0.006 | ** |
| Solanaceae.Cestrum.megalophyllum | 0.423 | 0.0268 | * |
| Group BC+BS+CLB |  |  |  |
| species | rpb.g | p.value | sig code |
| Cyatheaceae.Alsophila.erinacea | 0.42 | 0.0423 | * |

**Table 4B.** Indicator species for current ecological state (forest type)

| Group I |  |  |  |
| --- | --- | --- | --- |
| species | rpb.g | p.value | sig code |
| Boraginaceae.Cordia.colombiana | 0.661 | 0.0001 | *** |
| Melastomataceae.Meriania.tomentosa | 0.641 | 0.0001 | *** |
| Actinidaceae.Saurauia.sp.1 | 0.545 | 0.0003 | *** |
| Cyatheaceae.Cyathea.halonata | 0.458 | 0.0033 | ** |
| Fab.Caesalpinaceae.Senna.dariensis | 0.449 | 0.0046 | ** |
| Moraceae.Ficus.caldasiana | 0.42 | 0.0206 | * |
| Melastomataceae.Leandra.subseriata | 0.37 | 0.015 | * |
| Melastomataceae.Miconia.aff..brevithec | 0.368 | 0.027 | * |
| Piperaceae.Piper.fuliginosum | 0.356 | 0.05 | * |
| Staphyleaceae.Turpinia.occidentalis | 0.346 | 0.0426 | * |
| Group II |  |  |  |
| species | rpb.g | p.value | sig code |
| Cecropiaceae.Cecropia.andina | 0.811 | 0.0001 | *** |
| Cecropiaceae.Cecropia.sp.2 | 0.5 | 0.0009 | *** |
| Melastomataceae.sp.1 | 0.468 | 0.0051 | ** |
| Fab.Faboideae.Dussia.lehmannii | 0.334 | 0.0444 | * |
| Group III |  |  |  |
| species | rpb.g | p.value | sig code |
| Myristicaceae.Otoba.gordoniifolia | 0.524 | 0.0007 | *** |
| Cyatheaceae.Alsophila.erinacea | 0.515 | 0.0015 | ** |
| Lauraceae.Persea.aff..pseudofasciculata | 0.494 | 0.0022 | ** |
| Clusiaceae.Vismia.lauriformis | 0.44 | 0.0065 | ** |
| Melastomataceae.Conostegia.aff..centronioides | 0.418 | 0.0083 | ** |
| Arecaceae.Wettinia.aff..oxycarpa | 0.403 | 0.0115 | * |
| Lauraceae.Persea.pseudofasciculata | 0.386 | 0.0217 | * |
| Rubiaceae.Psychotria.paeonia | 0.373 | 0.0264 | * |
| Moraceae.Ficus.dulciaria | 0.354 | 0.03 | * |
| Group IV |  |  |  |
| species | rpb.g | p.value | sig code |
| Burseraceae.Dacryodes.cupularis | 0.754 | 0.0001 | *** |
| Burseraceae.Protium.ecuadorensis | 0.638 | 0.0001 | *** |
| Clusiaceae.Garcinia.macrophylla | 0.613 | 0.0001 | *** |
| Lauraceae.Beilschmiedia.aff..costaricensis | 0.448 | 0.0018 | ** |
| Melastomataceae.Conostegia.superba | 0.444 | 0.0024 | ** |
| Lauraceae.Ocotea.stenoneura | 0.369 | 0.0454 | * |
| Lecythidaceae.Gustavia.dodsonii | 0.362 | 0.029 | * |
| Styracaceae.Styrax.weberbaueri | 0.361 | 0.0408 | * |
| Moraceae.Pseudolmedia.rigida | 0.355 | 0.0263 | * |
| Group I+II |  |  |  |
| species | rpb.g | p.value | sig code |
| Cecropiaceae.Cecropia.ficifolia | 0.52 | 0.0005 | *** |
| Solanaceae.Solanum.lepidotum | 0.447 | 0.0055 | ** |
| Cecropiaceae.Cecropia.reticulata | 0.371 | 0.0244 | * |
| Verbenaceae.Aegiphila.alba | 0.348 | 0.0465 | * |
| Urticaceae.Urera.caracasana | 0.341 | 0.037 | * |
| Group I+IV |  |  |  |
| species | rpb.g | p.value | sig code |
| Lauraceae.Caryodaphnopsis.theobromifolia | 0.416 | 0.0092 | ** |
| Moraceae.Clarisia.biflora | 0.403 | 0.0157 | * |
| Group II+III |  |  |  |
| species | rpb.g | p.value | sig code |
| Annonaceae.Guatteria.megalophylla | 0.411 | 0.0133 | * |
| Moraceae.Ficus.cuatrecasana | 0.364 | 0.0299 | * |
| Group III+IV |  |  |  |
| species | rpb.g | p.value | sig code |
| Lauraceae.Aniba.aff..hostmanniana | 0.401 | 0.0138 | * |
| Capparaceae.Capparis.sp. | 0.385 | 0.0267 | * |
| Euphorbiaceae.Hieronyma.asperifolia | 0.383 | 0.0226 | * |
| Lecythidaceae.Eschweilera.integrifolia | 0.379 | 0.0199 | * |
| Lauraceae.Ocotea.insularis | 0.366 | 0.0295 | * |
| Rubiaceae.Faramea.oblongifolia | 0.362 | 0.0291 | * |
| Moraceae.Helicostylis.tovarensis | 0.346 | 0.0444 | * |

### Spatial analyses

13 Moran’s Eigenvector Maps variables were detected as correlated with changes in tree community (fig. 8, cumulated, adjusted R^2^ = 0.19, all MEMs p < 0.02 as components in forward model selection). Most were substantially correlated with 1-3 available environmental variables or a particular land-use/habitat history (table 5), with the exception of one highly localized MEM variable, MEM3. The most influential MEM variables in the model, MEM1 and MEM2 presented obvious geographic patterns. MEM1 (contributing R^2^ = 0.026 to cumulative R^2^, or 13.9% of explainable variance), indicates a general difference in tree communities between those of the northeastern and southwestern sides of the Los Cedros river (supp fig. 7). The second most influential MEM variable, MEM2, (contributing R^2^ = 0.026 to cumulative R^2^, or 13.6% of explainable variance) indicates a difference between the highest sites sampled in the study, which were situated along a ridge that runs ultimately to the highest points in the reserve, and lower elevations along this ridge system (supp. fig. 8). MEM8 also correlated with elevation, and with distance-to-nearest-stream, likely indicating a difference among “high and dry” sites and lower, wetter sites (supp. fig. 9).

**Figure 8.**
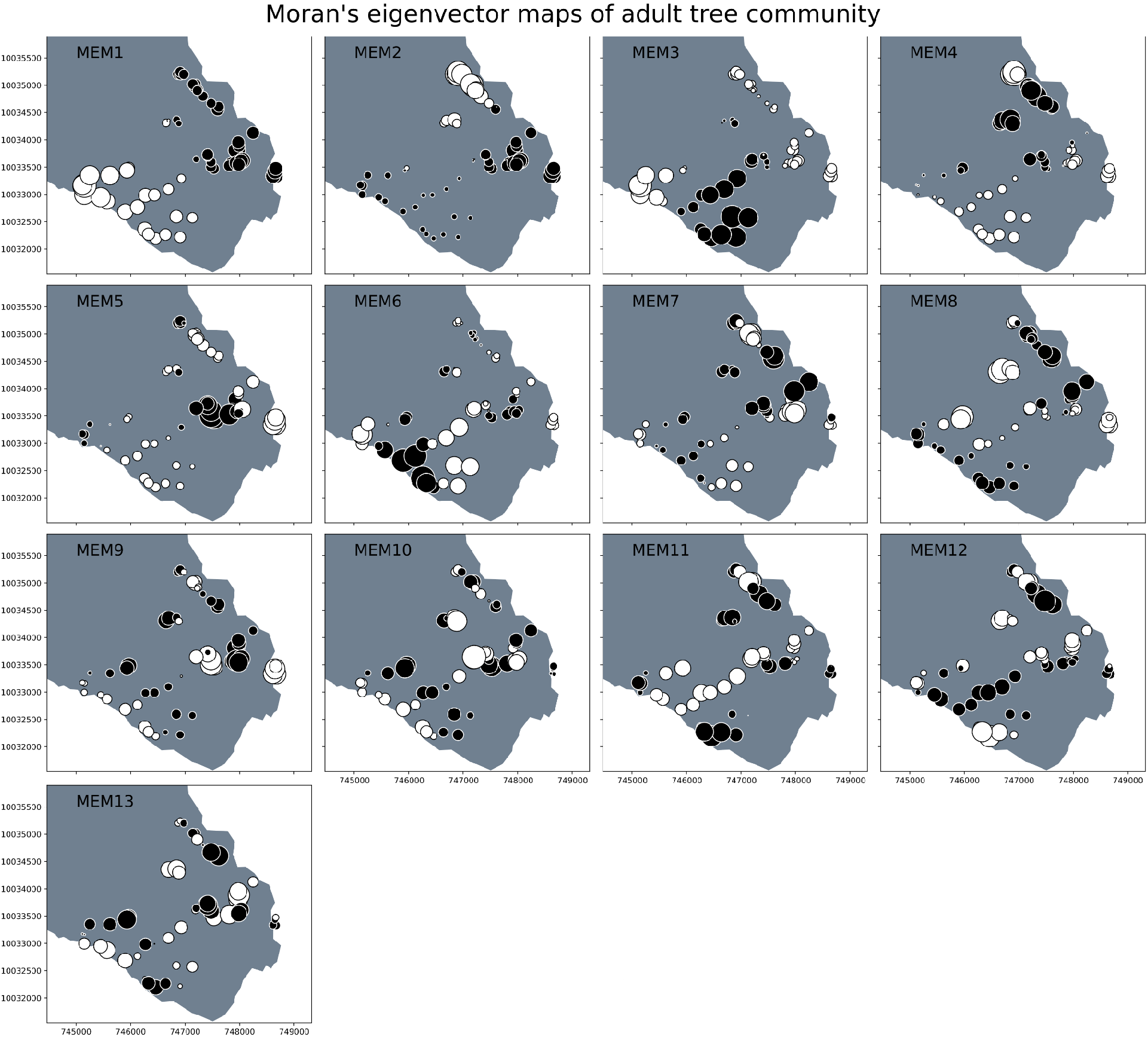

**Table 5.** Moran’s eigen vector maps and their statistically significant environmental correlations

|  | slope | DEM | aspect | exposeE | exposeN | toStream | BC | BS | CLB | RCA | RG |
| --- | --- | --- | --- | --- | --- | --- | --- | --- | --- | --- | --- |
| MEM1 |  | -0.403 |  |  |  |  |  |  |  | 0.287 |  |
| MEM2 | -0.344 | -0.636 |  |  |  |  |  |  |  | 0.308 |  |
| MEM3 |  |  |  |  |  |  |  |  |  |  |  |
| MEM4 |  |  |  |  | 0.349 |  |  |  |  |  |  |
| MEM5 |  |  |  |  |  |  |  |  |  | -0.503 |  |
| MEM6 |  |  |  |  |  | 0.306 |  |  |  |  |  |
| MEM7 |  |  |  |  |  |  |  |  |  |  | 0.385 |
| MEM8 |  | 0.404 |  |  |  | 0.460 |  |  |  |  |  |
| MEM9 |  |  |  |  |  |  |  |  |  | -0.451 |  |
| MEM10 |  |  |  |  |  | -0.340 |  |  |  |  |  |
| MEM11 |  |  |  |  |  | 0.282 |  |  |  |  |  |
| MEM12 |  |  |  |  |  |  |  |  |  |  | -0.410 |
| MEM13 |  |  |  |  |  |  | 0.345 |  |  |  |  |

### Juvenile communities

Juvenile communities did not cluster within the groups presented by adult communities, either in terms of historical land-use/habitat (fig. 9A) or current ecological state of adult trees (fig. 9B). Rather, all juvenile communities radiate into new dissimilarity space from their respective adult communities, possibly indicating recruitment of new tree species and/or new species combinations at these sites. Of the 148 species of juvenile tree species observed in the study, 110 species were observed as both juvenile and mature specimens, and an additional 38 species were observed only as juvenile specimens in the study.

**Figure 9A.**
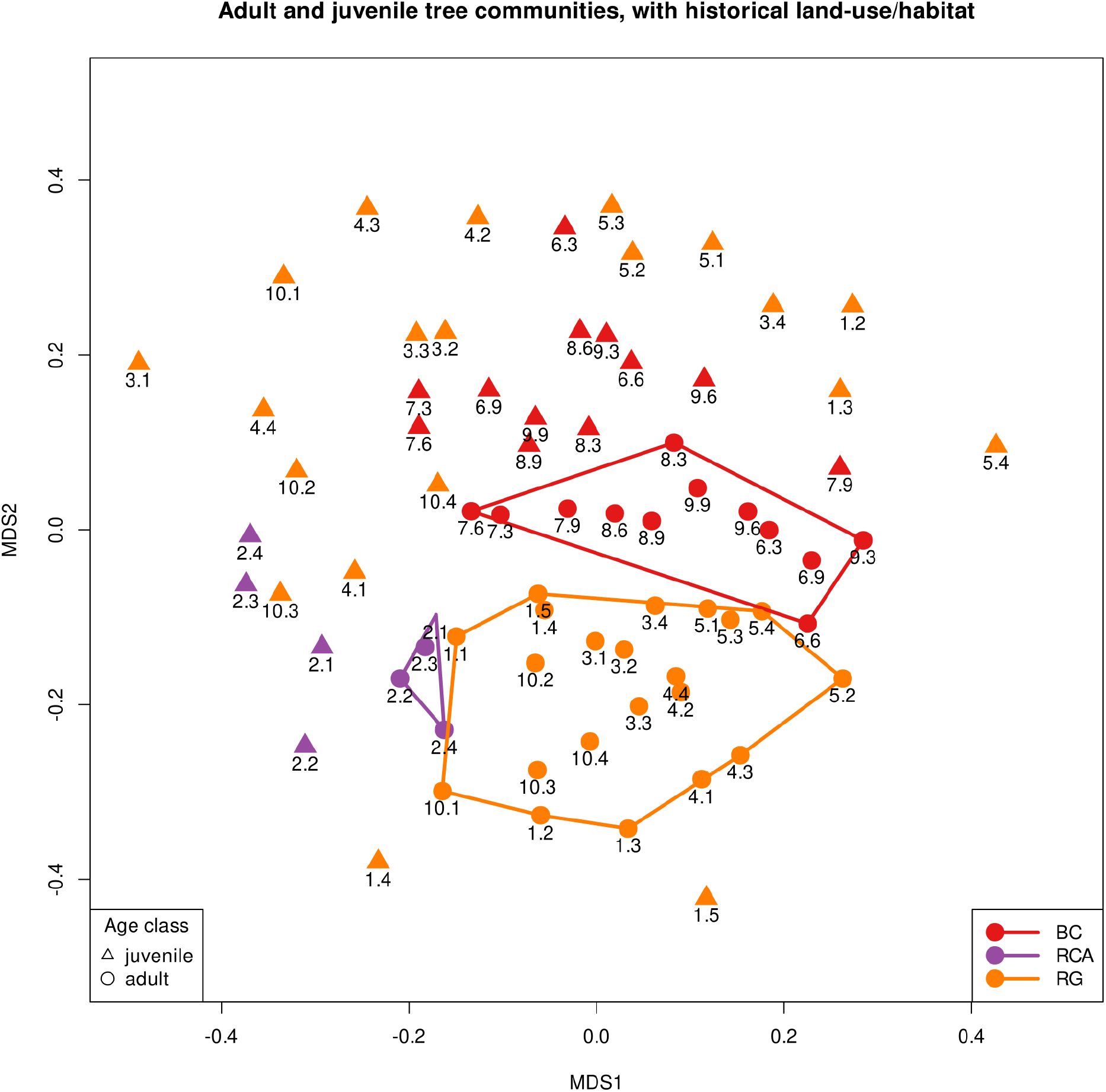

**Figure 9B.**
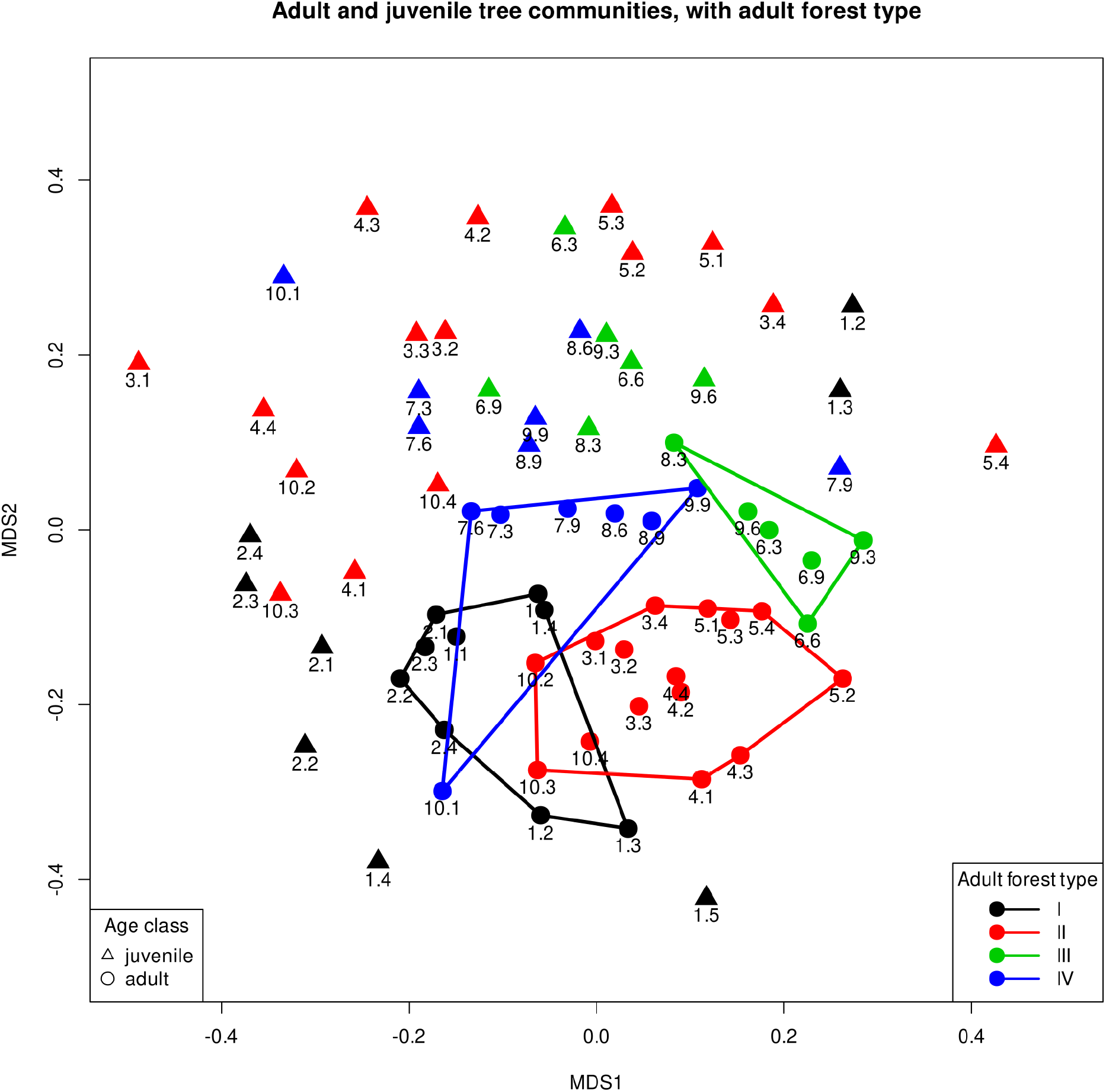

Juvenile tree communities change along the same axes as were observed for their respective adult tree communities. When categorized using historical land-use/habitat variable, RCA sites and related, similar RG sites are pulled in a negative direction along the NMS1 axis in fig. 9A, and non-anthropogenically disturbed sites (BC, BS, and CLB) are pulled in a positive direction along the NMS2 axis. When categorized by the current ecological state of their adult tree communities, the same pattern holds (fig. 9B): juvenile tree communities are different and outside of clusters formed by their respective adult communities, but change along the same axes as their adult communities. Two exceptions to this are observed: the juvenile tree communities of two type I forest sites have come to more closely resemble type III or forest (fig. 9B<u>, sites 1.2 and 1.3</u>). These sites have a land-use/habitat history of pasture-conversion (RG), and their adult tree communities cluster into a current ecological state of type I forest, the most anthropogenically disturbed group, often associated with RCA land-use history (fig. 5 and table.____). However their juvenile tree communities now resemble more “natural” forest types. Additionally, site 1.1 juvenile tree community was so unique as to be an outlier to the rest of the sites studied, and had to be removed for informative examination of the remaining sites (supp. fig. 10).

### Deforestation in the region

Between 1990 and 2018, forest cover in the Cotacachi canton was reduced from an estimated 87,967 ha of native forest-cover in year 1990 to 71,739 ha in year 2018, an 18% reduction of total forest from 1990 levels, mostly due to conversion to agricultural land (fig. 10). In terms of total land cover, this equates to a shift from approximately 52% of total Cotacachi canton land cover being native forest to 42% of total land cover as native forest. Forest cover in Los Cedros increased from an estimated 5,094 ha of native forest-cover in year 1990 to 5,210 ha in year 2018, an 2.3% addition of total forest from 1990 levels, due mostly to the reforestation of former pasture (fig. 11). Forest cover in Bosque Protector Chontal decreased from an estimated 6,920 ha of native forest-cover in year 1990 to 6,565 ha in year 2018, a 5% reduction of total forest from 1990 levels, due mostly to conversion of forest to agricultural land uses (fig. 11).

**Figure 10 A,B.**
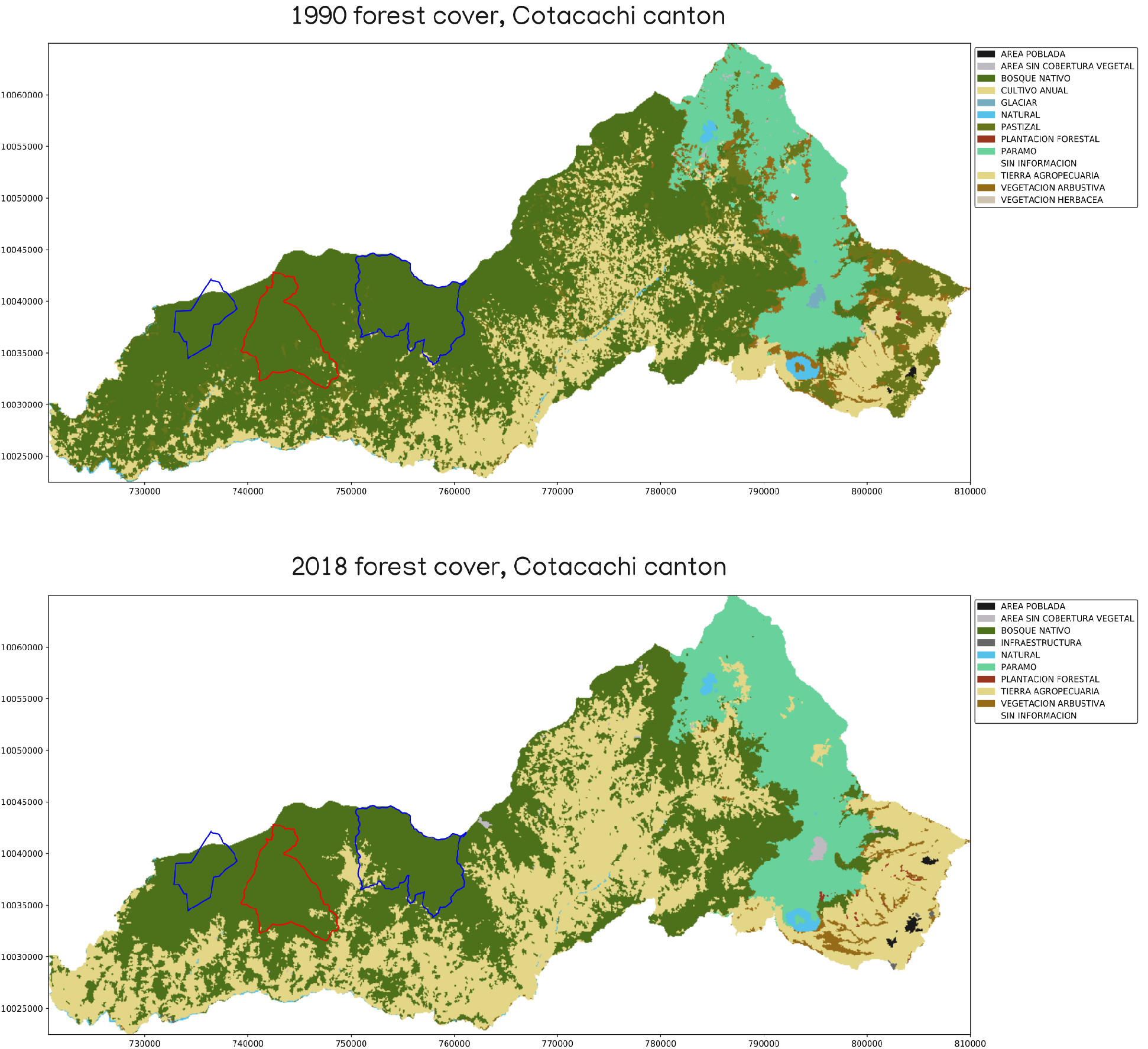

**Figure 11 A-D.**
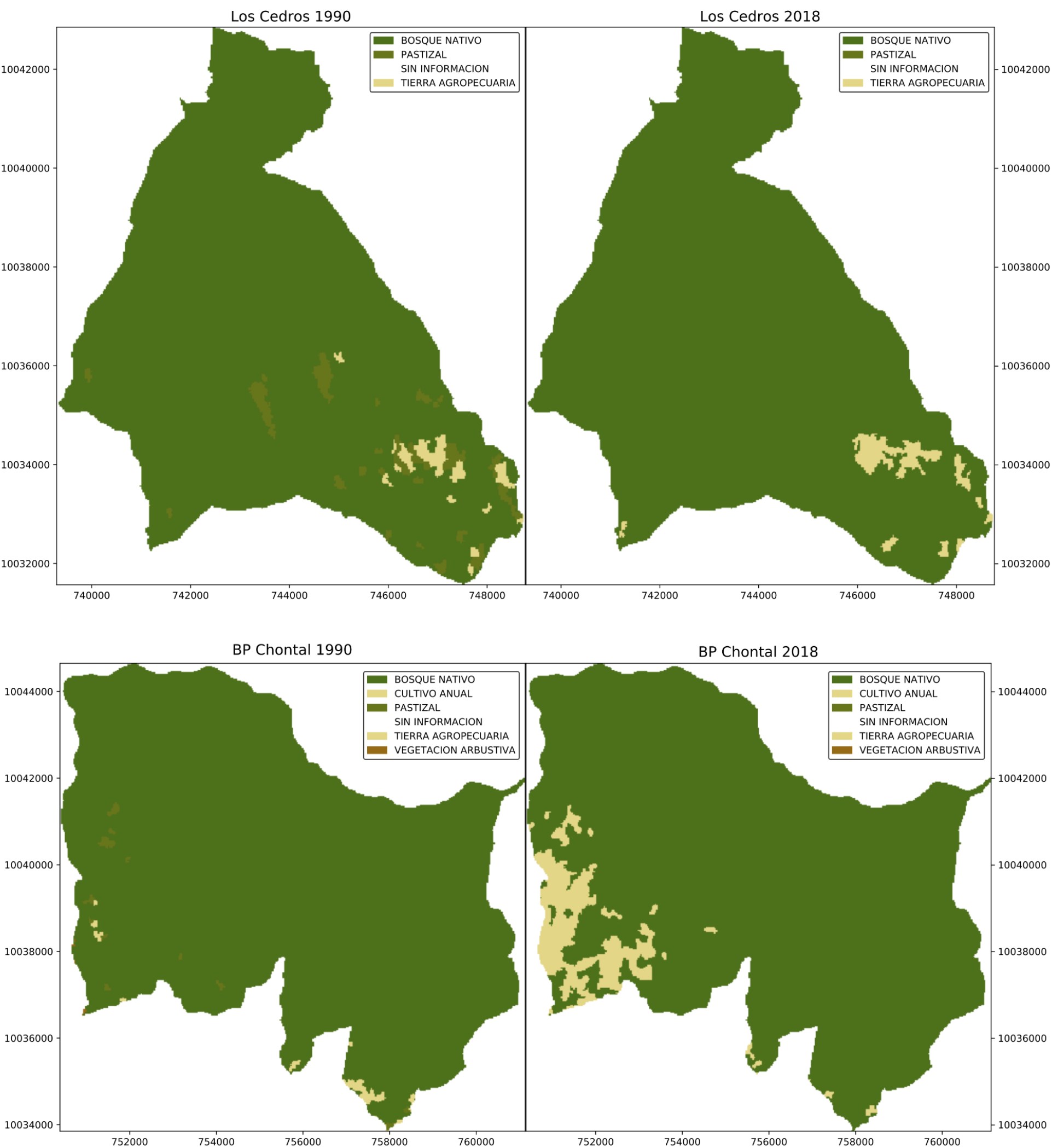

Unlike earlier datasets, the 2018 MAE land cover dataset does not include pasture separately from other forms of agricultural land use, so it is difficult to quantify which types of agricultural land-use that are most commonly replacing forest in the region. However, personal communication and visitation with the surrounding communities, it is the authors assessment that much of the recent deforestation is undertaken to create pasture for cattle grazing.

## Discussion

### i. Stable states in the Andean cloud forest

In this study we examined 61 forest sites with known histories of anthropogenic disturbance or natural, gap-forming disturbance, and examined this history as a predictor of current ecological state. We can categorize the southern region of the Los Cedros reserve into four forest types, based on adult tree communities (fig. 4), and well predict these ecological states from past land-use/habitat-type and elevation. Sites that have no history of anthropogenic disturbance group readily into either forest types III or IV (fig. 4, fig. 5). These sites are characterized by endemic, forest-dependent tree species (table 4B). The difference among these “natural” forest types III and IV are strongly predicted by elevation (fig. 7, fig. 6, supp. fig. 5), exhibiting elevation-dependent ecological zonation often observed in montane tropical forests (Grubb, 1977; Leigh, 1975). Natural gap-forming disturbances do not change the species compositions of these sites to make them significantly different from other natural forest sites (fig. 5). As such, these natural forest types may represent stable equilibria with basins of attraction that give each some resilience to gap-forming disturbance (fig. 12).

**Figure 12.**
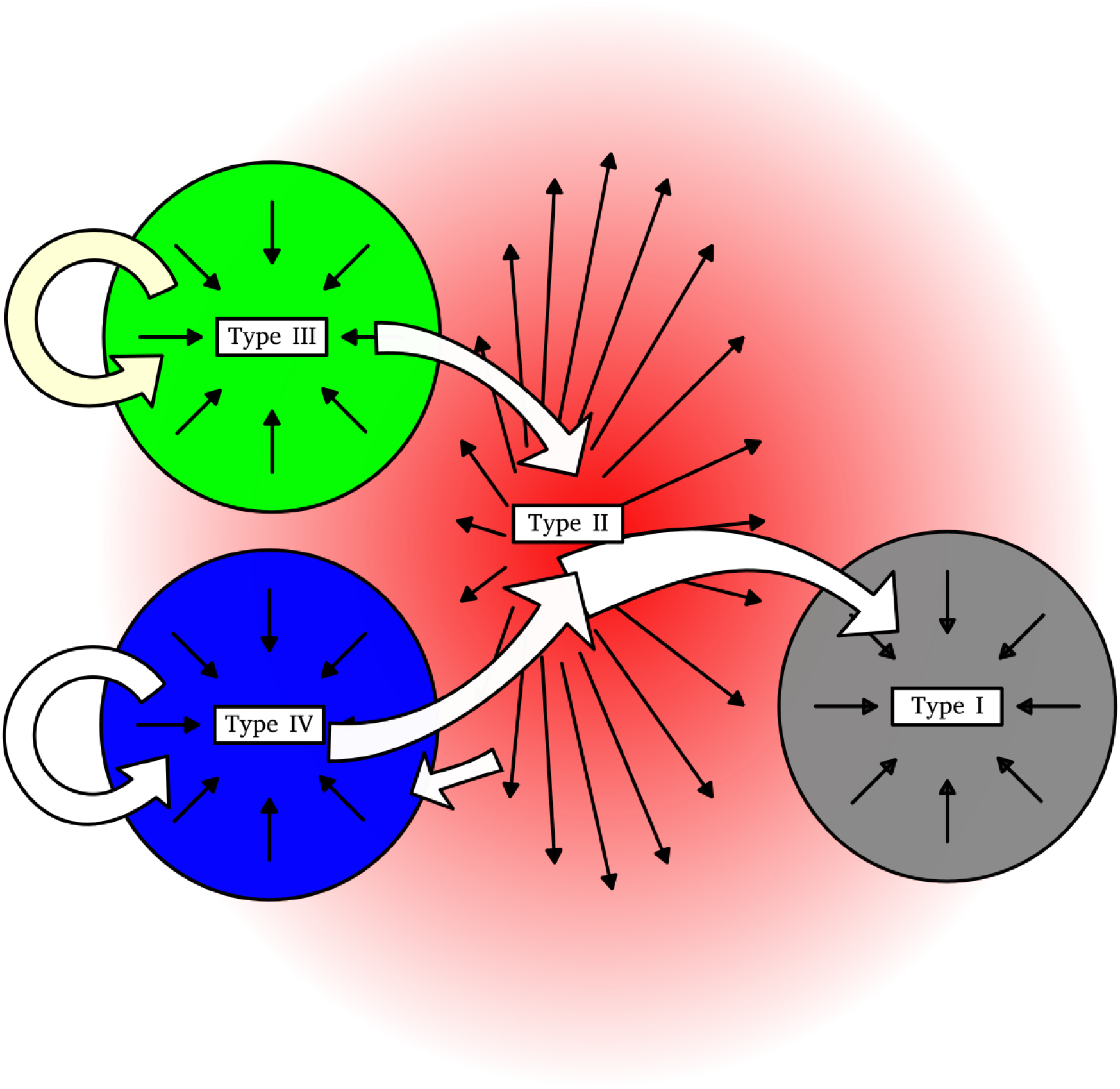

There are indications that sites that have experienced intermediate disturbance (conversion to pasture, followed by reforestation) have entered a less stable, more pluripotent ecological state. Nearly all high and low elevation sites that were converted to pasture prior to reforestation (land-use/habitat type “RG”) then developed into a single community type, (“forest type II”). This suggests a homogenizing effect on tree communities from this kind of disturbance, and a common successional response by the forest to after conversion to pasture and abandonment, regardless of elevation. In ordinations of community similarity (fig. 5 and supp fig. 3), Type II forest sites are situated in an intermediate position between all other states. It is therefore possible that the Type II forest type represents a low-sloped, convex unstable equilibrium that can sometimes allow sites to return to a natural forest state. In one case, we observe a site that has undergone anthropogenic disturbance and subsequently develops into a natural forest type (site 10.1, circled in fig. 4, fig. 5). Additionally, in the case of two pastured (RG) sites, their adult communities resemble the highly affected Type I forest, but their juvenile communities currently more closely resemble the “natural”type III forest (fig. 9, sites 1.2 and 1.3). In a different successional direction, several pastured (RG) sites developed into the novel Type I forest, the same forest type into which all intensive-agriculture sites developed (fig. 5).

The existence of this indeterminate, intermediate ecological state, forest type II, supports the hypothesis that primary cloud forest has some capacity to “repair” highly modified agricultural sites, but also indicates that this is not a certain outcome. This post-pasture successional trajectory appears to be very distinct from that which follows natural, gap-forming disturbances, and may be intermediate to severe agricultural conversions (cane production, followed by reforestation), which tended to drive sites to another state, forest Type I. The adult tree communities of sites which experienced intensive agriculture all developed into forest type I, suggesting that forest type I may be another stable equilibrium or novel ecosystem type (Hobbs et al., 2009). Additionally, all sites that were observed to be in Type I state are low-elevation sites, sites that presumably would have otherwise existed in a Type IV state, suggesting particular vulnerability of these lower-elevation forests to anthropogenic disturbance. However, all type I sites are also co-localized to the same area of the reserve (fig. 6), making it difficult to generalize this pattern to the rest of the study area.

Several forces may be at work in creating these equilibria we observed. Type III and type IV forests (“natural forest” types), gap-forming incidents appear to keep these sites within their basins of attraction, and the breakdown of these negative-feedbacks and ecological conditions that are likely responsible for shifts into forest type I equilibria:

#### Soil structure

While soil structure data are not available for this survey, researchers at field sites noted that sites that had undergone intensive, *cañaveral* agricultural use (“RCA” sites) had more compacted soils. Agricultural use can incur long-lasting legacy effects on soil structural and chemical characteristics, even after abandonment and reforestation (Sohng et al., 2017; Zhang et al., 2019). Deforestation has been shown to cause long-term changes in soil physical structure and microbial activity in soils, especially among plant-symbiotic microbes (Hartmann et al., 2012, 2014; Wang et al., 2018).

#### Soil seed bank depletion

In addition to soil compaction, several seasons of indiscriminate grazing or sugar cane culture probably greatly depleted the residual seed bank of forest plants (Bart & Davenport, 2015; K. R. Young et al., 1987). When soil-seed banks have been exhausted, recolonization of sites by forest plants must occur through dispersal of seeds to the reforesting site.

#### Large seeded plants and animal dispersal

Reserva Los Cedros hosts three species of primates: the brown-headed spider monkey (*Ateles fusciceps fusciceps*), the white-headed capuchin, (*Cebus capucinus*), and the mantled howler monkey, (*Alouatta palliata*), in addition to numerous other frugivorous birds and mammals (Roy et al., 2018). These animals likely play an essential role in closing gaps after disturbance. In recent natural gaps (‘CLB’), we observed a single indicator species - *Endlicheria* sp. (table 4A). Some *Endolicheria* species have been suggested elsewhere to be primate dispersed (Bufalo et al., 2016), supplemental materials). Lower elevation natural forests (forest type IV) were characterized by copal trees (*Protium* and *Dacryodes* spp., table 4B), well known locally as primate food sources (Morelos-Juárez et al., 2015). In general, Type III and IV forest types were generally characterized by plants with larger seeds or fruits, often primate or otherwise vertebrate-dispersed, such as those in Lecythidaceae and Lauraceae, and *Garcinia* (table 4B). This prevalence of trees with larger seeds/fruits is typical of tropical forests, where most trees produce animal-vectored seeds (Howe & Smallwood, 1982). Absent active transport of seeds by animals, colonization of pastures by forest trees is expected to be very slow (Cubiña & Aide, 2001), and to heavily favor wind dispersed seeds (Holl, 1999). Many ruderal species, conversely, are wind-dispersed (Grime, 1977) and are likely to readily colonize abandoned field sites, especially when aided by human vectors (Planchuelo et al., 2020). Given Los Cedros’ relatively low elevation among montane tropical forests, it is not surprising that it’s forests are heavily populated with large seeded species that likely rely on primate dispersal (Chapman et al., 2016).

#### Relative fluxes of local vs. exotic plant types

Most of the sites examined here are embedded in a landscape of primary forest, and may have been able to recruit seeds from forest dependent species even if they experienced seed bank depletion (Willson & Crome, 1989), while still insulated from outside seed sources. Approximately half of type I forest sites, however, were located on the edge of los Cedros, abutting neighboring farmland, probably allowing for extensive input of small seeded pioneer tree species such as *Saurauia* spp. The remaining type I sites were located along mule trails that supply the reserve, and were also therefore presumably repeatedly exposed to seeds from outside locations.

The above mechanisms are primarily “gap-filling” mechanisms, and may fail in the face of extensive anthropogenic disturbances such as habitat fragmentation and loss (for example, Cramer et al. (2007)). It is probably important to note how dependent these processes may be on large animal dispersers, and especially on primates, as primate populations are in decline in the region (Peck et al., 2011; Roy et al., 2018). The importance of seed-dispersal services by large mammals and spider monkeys in particular can be appreciated by anticipating their loss: Peres et al. (2016) have predicted substantial loss of forest biomass in forests if large mammal populations are reduced, especially spider monkeys and tapirs. A similar positive feedback may be possible with the loss of forest-dependent birds due to deforestation. Frugivorous, forest-dwelling birds provide unique seed-dispersal service to forest trees but are highly interdependent with their habitat; their decline in diversity has been directly linked to vegetative diversity loss and deforestation (Morante-Filho et al., 2018).

Other complex interactions may also be important: the presence of *Cecropia* spp. as indicator species in forest types with intermediate disturbance (forest type II), and appears less often in the novel type I forest. In this study they were also not observed as extremely prevalent in recent natural gaps. This is possibly important as *Cecropia* trees occupy a somewhat unique position of being an early-successional species while also producing large fruits and canopy structure characteristics useful to both primates and frugivorous birds (Morelos-Juárez et al., 2015; Navarro et al., 2018; Skutch, 1945). *Cecropia* trees may attract primate dispersers and forest-dependent birds back to a disturbed site, and therefore bring with them the heavier seeds of other forest-dependent, heavy-seeded species. At Los Cedros, the smallest primate dispersers, capuchin monkeys, are regularly observed in *Cecropia* trees. *Cecropia* spp. may therefore act as another gap-repairing negative feedback mechanism, one that plays out in cases of forest gaps of larger size or severity than the smaller natural gaps examined here. Thus the success or failure of *Cecropia* may play a deciding role that allows sites to return from the ecological state we have here observed as forest type II (resulting from intermediate disturbance), and potentially back to primary-like forest states. Note again, however, that this mechanism would depend upon the presence of large animal vectors, especially primates.

Given the relatively short period of time since disturbance prior to surveys (13-18 years), it is also very difficult to confidently project the stability of the here-proposed candidate equilibria to larger time scales, especially in times of deep ecosystem change due to climate change. Sites were surveyed in 2005, so an additional 16 years have passed since the collection of these data. Indeed, Loughlin et al. (2018) suggest that tropical Andean cloud forest in a nearby ecoregion may have a single long term ecological equilibrium. However, conditions in the Andes are changing dramatically from those that presumably sustained the long-term resiliency observed by Loughlin. Thus the candidate alternative states observed here should not be disregarded, they may mark the beginning of novel forest types in the region (Hobbs et al., 2009). For the moment, visual inspection of these sites in the present day confirms that unique tree communities continue to exist at Type I forest sites compared to surrounding older forest. Updated systematic surveys are now necessary to investigate the stability of the ecosystem types suggested here.

### ii. Juvenile tree community

It is difficult to assign cause to the lack of structure observed in juvenile tree communities at Los Cedros (fig. 9). Much of the noise in these juvenile communities presumably stems from the incomplete filtering of young plants at each site (Baldeck et al., 2013; Ledo et al., 2015) - the environmental pressures that have shaped the adult tree community at each site likely had yet to totally act on the juvenile trees at time of sampling. However, it is also inevitable that the same global environmental changes that are acting on other forests throughout the world (Allen et al., 2010; Seidl et al., 2017), including cloud forests (Foster, 2001), are at play in the ancient forests of Los Cedros, and are contributing to the reorganization of future tree communities, perhaps uncoupling ancient species associations. The disturbance regime under study (conversion to agriculture, followed by reforestation) also introduced new conditions and plant species to Los Cedros. Thus in these juvenile trees we may also be observing two sources of disturbance that could shift the forests of los Cedros out of the basins of attraction of the primary forest state, of very different scales: local land use change, and global climate change. In the terminology of Beisner et al. (2003), the former may still be considered a state variable change from which it is sometimes within the capacity of the cloud forest to rebound and recover to a primary forest state. The latter, however, is a deep shift in the parameters of the Andean cloud forest ecosystem, which will no doubt change the shape of the possible (Clerici et al., 2019; Feeley et al., 2011; Rehm & Feeley, 2015). It is not unreasonable to expect interactions between these two fundamental sources of ecological disturbance (Clerici et al., 2019; Pimm et al., 2014; Stork, 2010; Tovar et al., 2013).

### iii. Beta-diversity and spatial heterogeneity in the Andean cloud forest

We examined patterns of distance decay in the tree communities of the southern region of los Cedros. When community turnover is modeled as a function of euclidean distance, a model in the form of an asymptote function that fits well to observed patterns of community-turnover. Our asymptote model suggests turnover in short distance, reporting half of maximum dissimilarity is reached in just ∼150m, and a mean BC > 0.8 in comparisons with distances larger than 600m (fig. 2). When small watersheds are used as the basic spatial unit, rather than Euclidean distance, most of the decay in tree community similarity occurs with the first crossing over to a neighboring watershed (fig. 3). Following this, mean dissimilarity among sites increases slightly but remains uniformly high, and further comparisons are not statistically significantly different, meaning comparisons between sites five watersheds away were not on average more or less similar than comparisons of sites that were only two, three or four watersheds apart, because so much change in community composition has already occurred just within the first watershed crossing. This is supportive of the colloquial understanding that in the Andes, each small drainage can host an almost entirely distinctive community from its neighbors.

This may also give more insight into the processes that create the fine-scale of endemism often observed in Andean species. We hypothesized that this fine-scale, watershed-based community turnover is due to high dispersal limitation and high micro-site variability that result from the complex, dramatic topography of the Andes. This microsite variation is visible to some degree in our Moran’s eigenvector maps and their environmental correlations (fig. 8, supp. figs 7-9, table 5): we see that environmental variables vary on scales that can be well understood in terms of change within and among small watersheds, and that these spatio-environmental patterns may explain up to 19% of variance in the plant community. This fraction may represent much of the environmental filtering that is occurring within the tree community at los Cedros. Dispersal limitation is harder to test, but the predominance of large-seeded species observed in our ancient forest sites suggests that many important tree species are dispersal limited to highly local scales and dependent on large animal dispersers to overcome this limitation. This preponderance of heavy-seeded species also might be well approximated by a symmetric dispersal-limited neutral model, a model which can generate significant small-scale spatial patterning in communities even without considering additional microsite environmental conditions (Hubbell, 2008).

### iv. Conservation value of Los Cedros

Since the sounding of alarm by Gentry, Myers et al., and numerous others (Gentry, 1992; Myers, 1988; Myers et al., 2000), there has been much concern about the future of the Chocó and the tropical Andean Biodiversity hotspot by conservation groups. However, there is little evidence that this call for conservation has resulted in sufficient meaningful change for the region of Los Cedros - quite the opposite. Instead, during this time frame, Cotacachi Canton has lost significant forest cover (fig. 10), as has Ecuador generally (Dodson & Gentry, 1991; Mosandl et al., 2008; Tapia-Armijos et al., 2015). In stark contrast, Los Cedros has well withstood the traditional pressures of timbering and settlement, as evidenced by its increase of forest cover during a time of net forest loss in the Cotacachi Canton. In addition to the historical habitat conversion from timber extraction and settlement, the tropical Andean Biodiversity hotspot has now found itself in a new center of metals mining exploration (Guayasamin et al., 2021; Roy et al., 2018; Vallejo Galárraga & Freslon, 2017), a new and entirely different extractive pressure on the region. Los Cedros itself is targeted by a junior mining partner (cornerstoneresources.com). Los Cedros has responded with a legal challenge that has reached Ecuador’s highest courts, with implications for all Bosque Protectores in Ecuador (Guayasamin et al., 2021). Due to its proactive defensive legal efforts, Los Cedros’ conservation effect is thus amplified even beyond the unusually effective physical protection of its forests.

Going forward, however, conservation of primary forest reserves such as Los Cedros will face novel challenges. It is widely understood that the high endemism of the Andean biodiversity hotspot makes it both a conservation priority and an especially difficult conservation challenge. It is less widely appreciated that forests, and especially primary forests, pose a major additional challenge to conservation. In most cases, we have only a crude understanding of the numerous ecological interactions required to maintain a primary ecosystem in its current state of biodiversity and ecosystem services, otherwise known as the ecological complexity of an ecosystem (Filotas et al., 2014; Messier et al., 2015). Ancient ecosystems are unique in large part due to their emergent complexity, and quantifying this complexity is usually difficult, requiring composite measures of numerous indices of forest physical structure, plant community, and other ecological properties (Loehle, 2004; McElhinny et al., 2005). In complex ecosystems, the loss of a species is multiplied by disruption of its interactions with other species, such as with trophic cascades (Ripple et al., 2016). The number of possible interactions among species undergoes quadratic growth as biodiversity increases linearly, so the immense biodiversity of the tropics presents a particular challenge to understanding the local ecological complexity. Primary forest fragments are also presumably subject to the well-known vulnerabilities of “island” - or insular - habitats (MacArthur & Wilson, 2001; Ross et al., 2002), namely the stochastic extinction of species without any nearby “mainland” to replenish populations. This is in addition to the uniquely dynamic, weather-dependent boundaries which make all cloud forests insular systems, primary or otherwise, and that may make them very vulnerable to shrinkage from climate change (Foster, 2001; Gentry, 1992; Rehm & Feeley, 2015). Thus in insular, highly interdependent ancient ecosystems such as primary cloud forest, these stochastic species extinctions may have greater cascading impacts than in simpler ecosystems. Even if completely protected from external changes, primary forest fragments such as los Cedros may ecologically simplify or “decay” with time if they are not sufficiently understood, buffered and interconnected. As such, the regional resiliency noted by Loughlin et al. (Loughlin et al., 2018) may be subject to novel vulnerabilities that were not so urgent historically. Michaels et al. (2020) have suggested that reforestation efforts should first attempt to diagnose the ecological conditions and feedbacks that allow a basin of attraction for a desired stable state to persist, and also find the minimum “nucleus” area needed for those processes to proceed and grow. They and others, such as some proponents of rewilding (Fernández et al., 2017), suggest seeding landscapes with species consortiums intended to recreate ecological complexity as much as possible. Los Cedros is far from needing rewilding, but conservation efforts around Los Cedros and other ancient tropical forests will be greatly enriched by such an awareness of protecting and nurturing ecological complexity, beyond simple physical protection of reserve borders.

Given the state of fragmentation and loss of forest in the region, and the poor understanding of its numerous pollination and diaspore dispersal networks, especially the threatened state and habitat requirements of the primate dispersers of los Cedros (Peck et al., 2011), Los Cedros may actually be at or close to threshold stable patch size for maintaining its level of biodiversity. Los Cedros may therefore face an important juncture, with a choice of growing or subsiding. Further reduction or anthropogenic disturbance of cloud forest area risks non-linear, catastrophic changes. Additionally, the fragments of Andean primary cloud forest habitat that persist to date are rare and small enough that there is no economic justification for eliminating or heavily modifying them. Simply as a matter of scale, any economic benefit from the degradation of these forest fragments will not be large enough to justify their loss as reserves of genetic information, providers of ecosystem services, and protectors of species diversity. Other methods of economic development and poverty reduction are available (Balmford et al., 2002; Kocian et al., 2011). However, great opportunity for the future exists in Los Cedros: the forest of Los Cedros need not be seen as a delicate ecological island in a storm, nor is its protection an academic or moral exercise, or a desperate rear-guard conservation strategy. It is instead useful to think in the sense of Michaels (2020), supported by the long-term observations of Loughlin et al. (2018), and see the Los Cedros forest as a nucleus of primary cloud forest that has prospered for thousands of years, that has weathered the recent waves of deforestation, and that now stands ready to help reseed the new forests of the northern Andes.

## Acknowledgements

The work would not have been possible without Jose DeCoux and his dedication to the conservation of Reserva Los Cedros. Several field assistants, volunteers and guides associated with Los Cedros helped with data collection: Gila Roder, Martín Obando, Manuel Moreno, Danny Cumba, Homero Sánchez, Fausto Lomas and Víctor Lomas.

## Supplemental Figures

**Supplemental figure 1.**
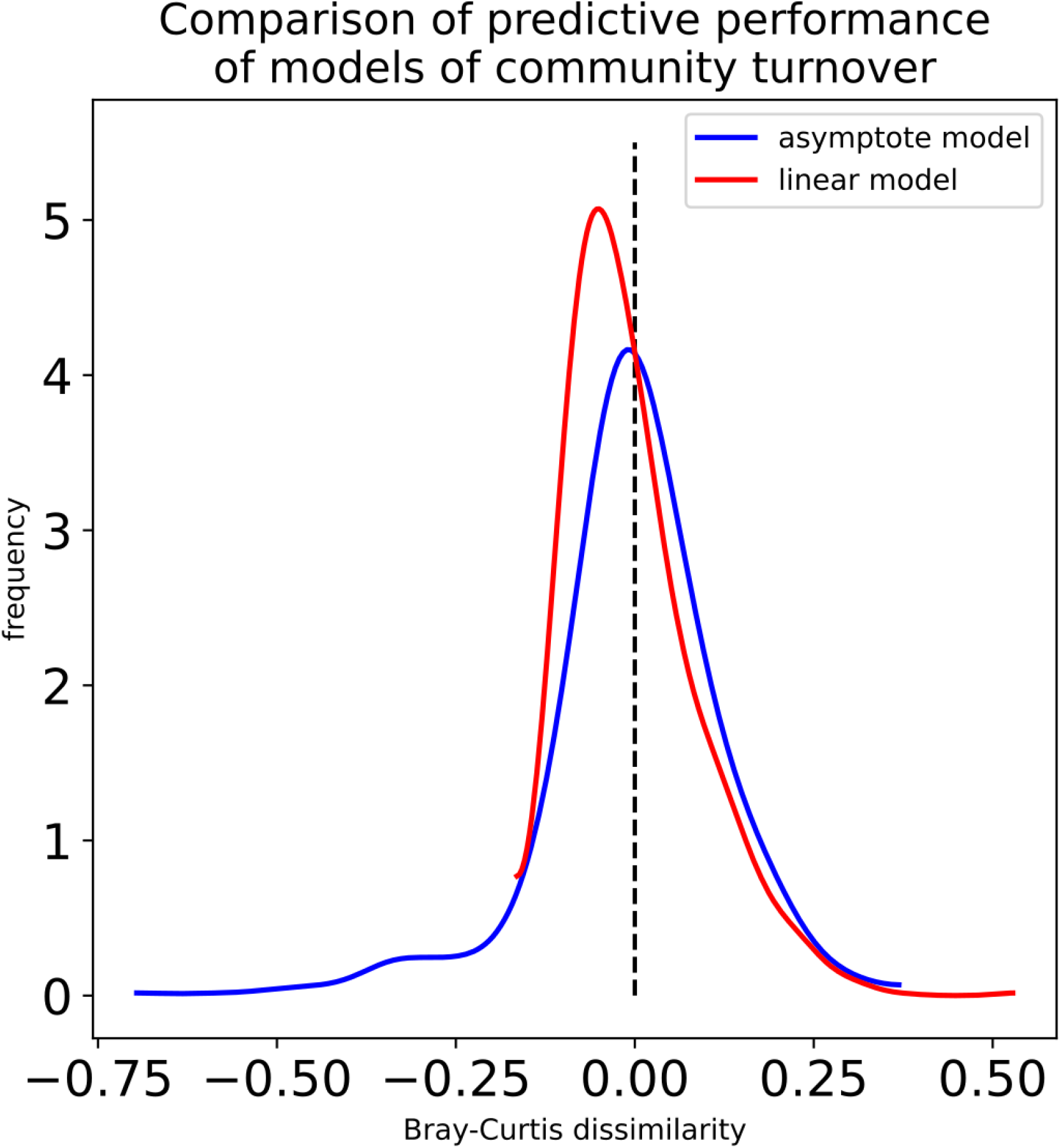

**Supplemental figure 2.**
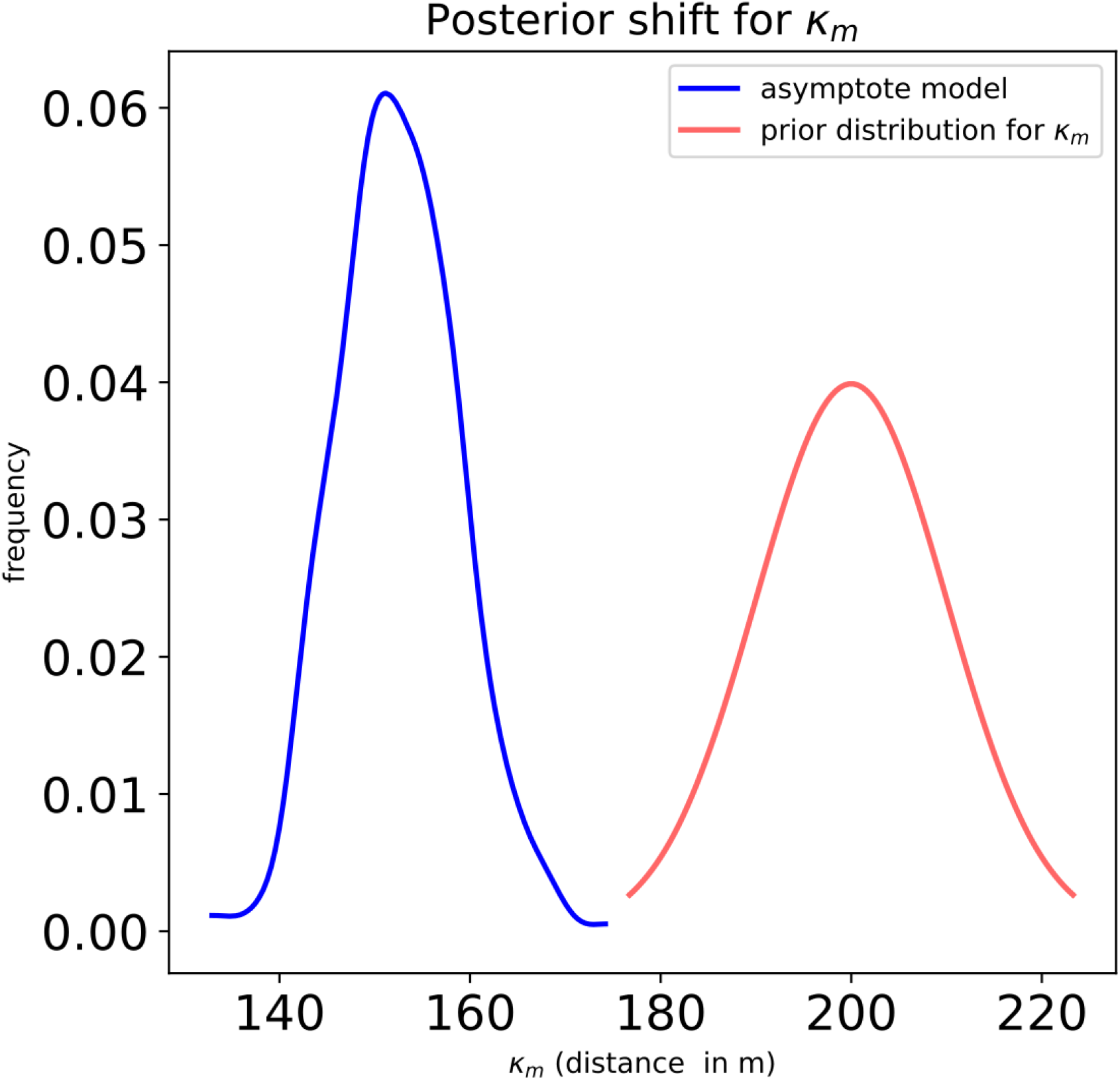

**Supplemental figure 3.**
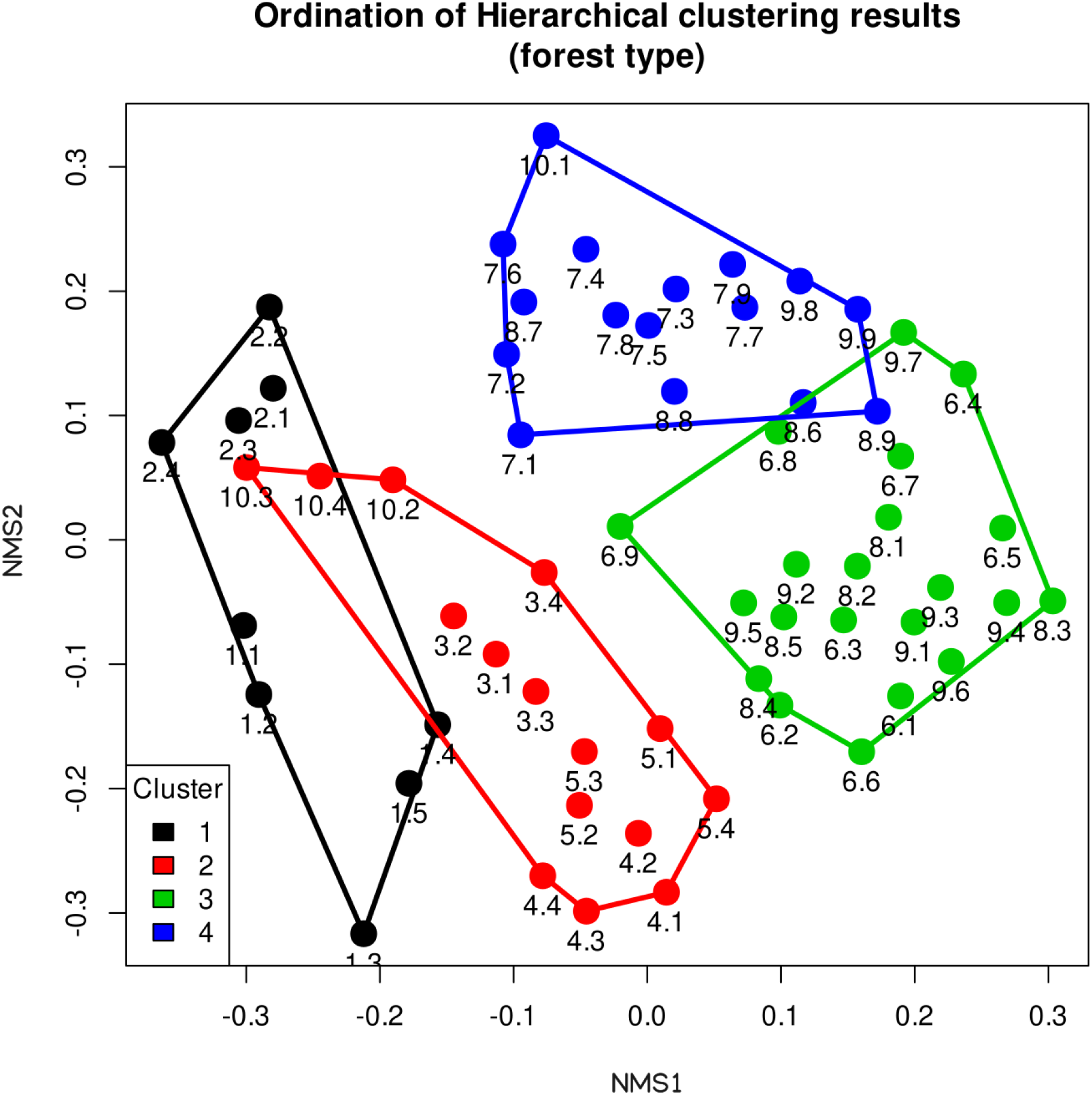

**Supplemental figure 4.**
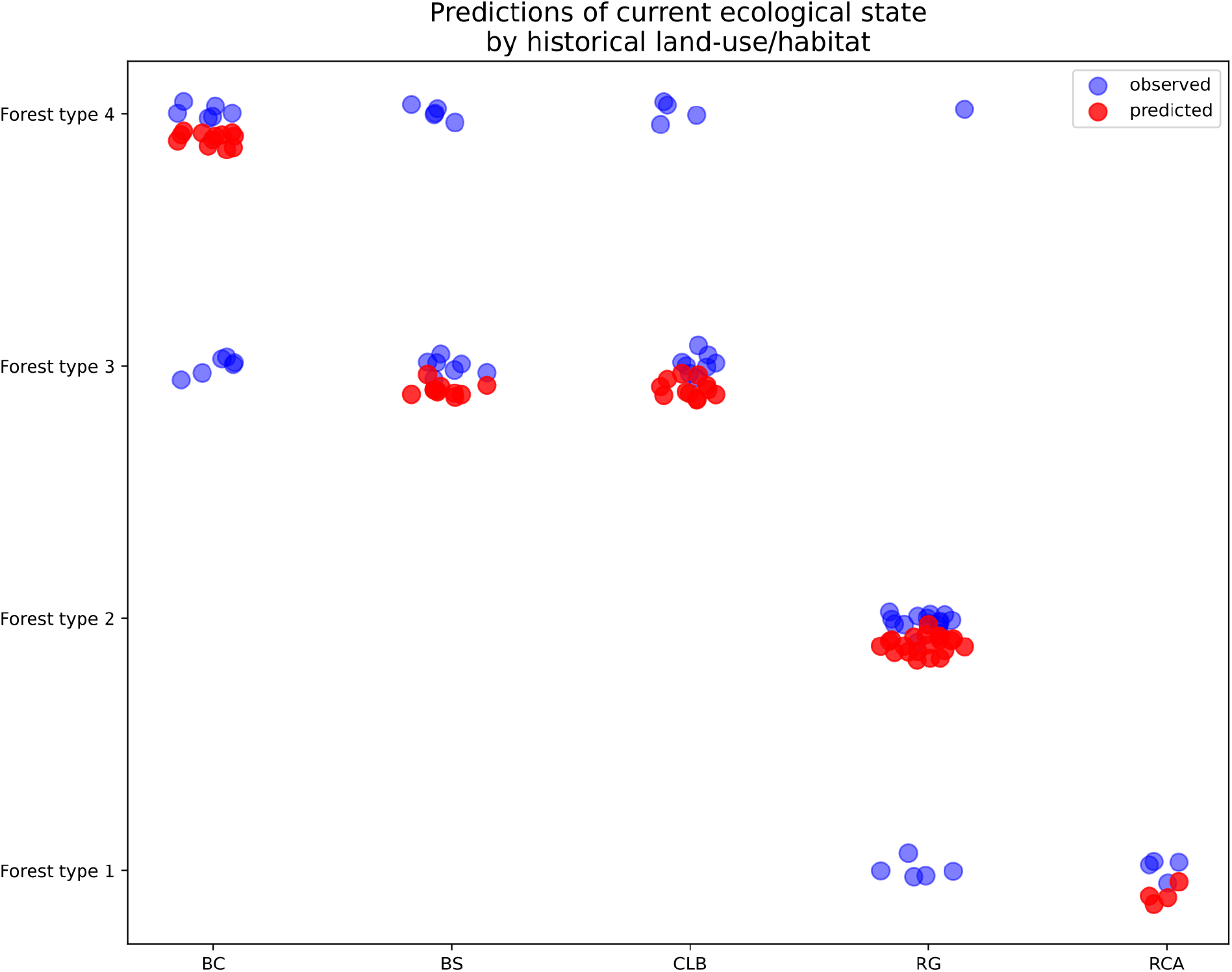

**Supplemental figure 5.**
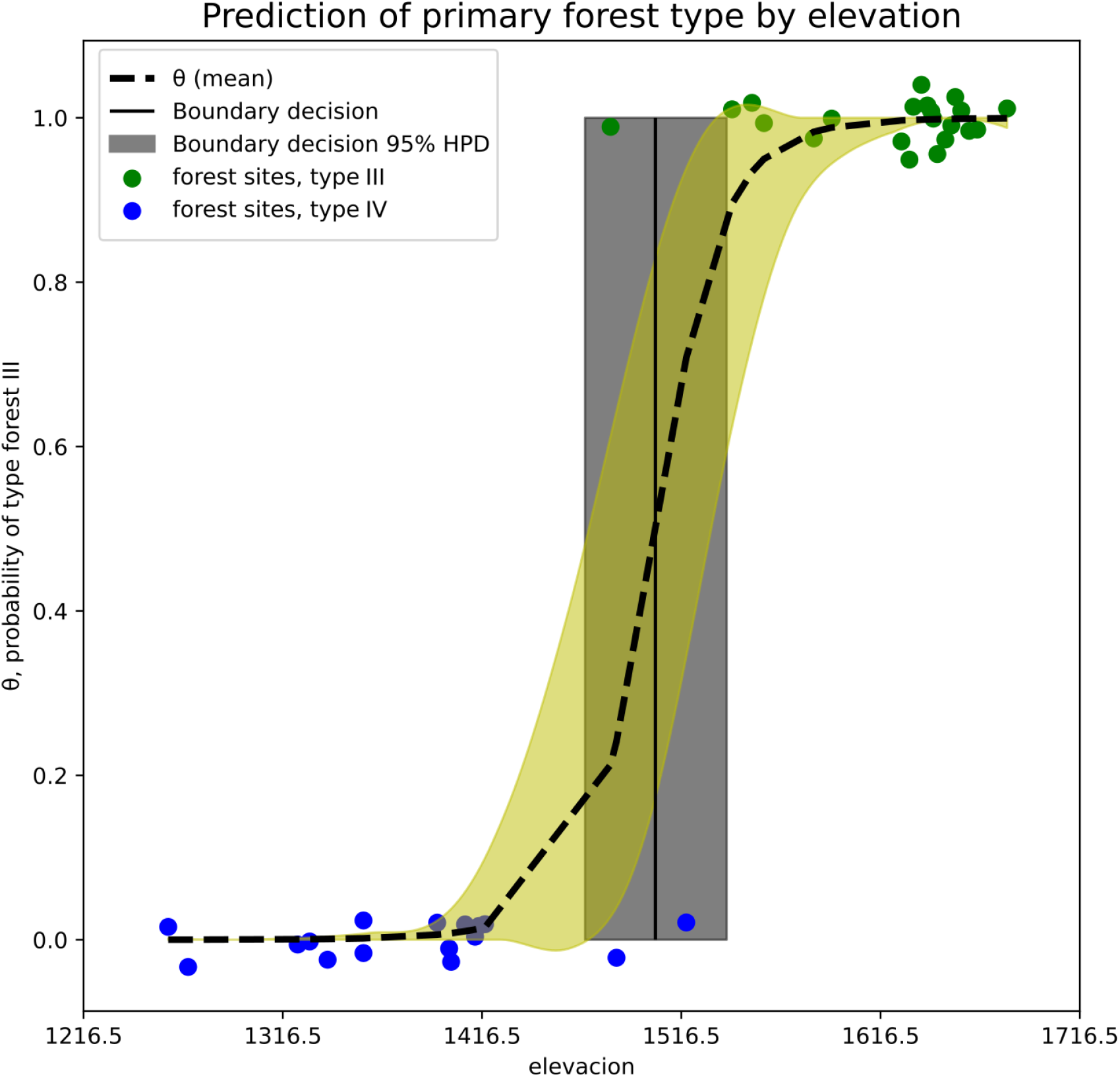

**Supplemental figure 6.**
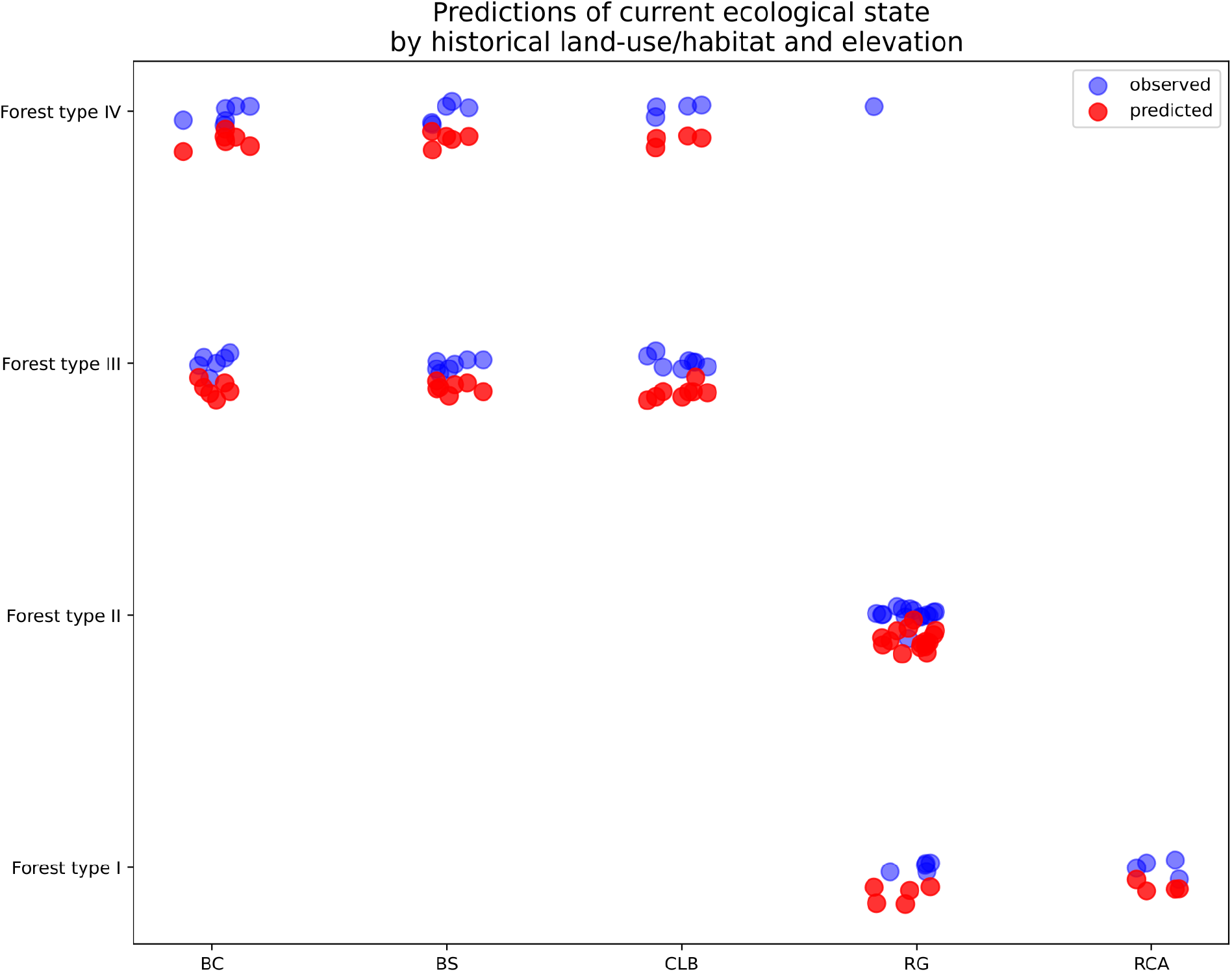

**Supplemental figure 7.**
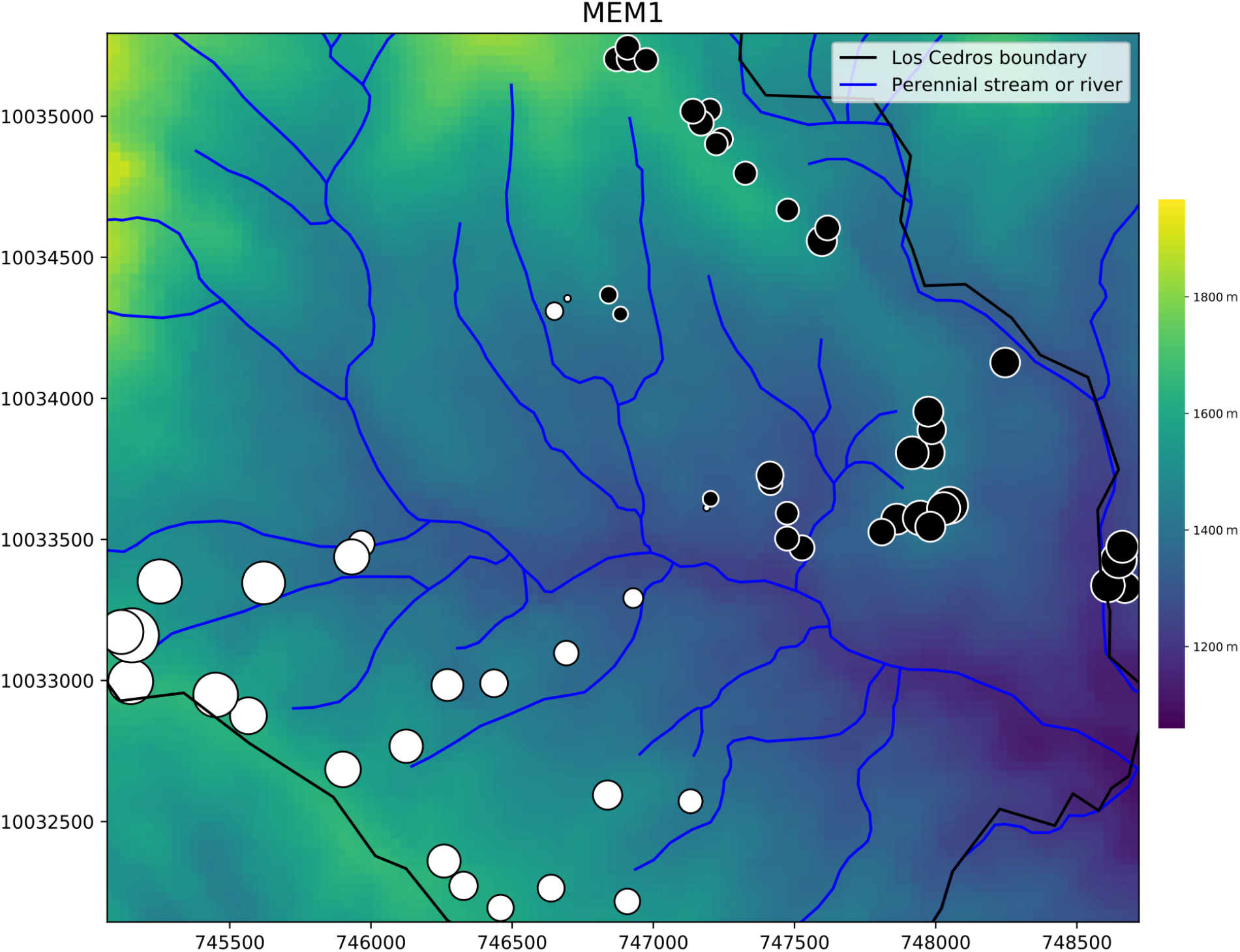

**Supplemental figure 8.**
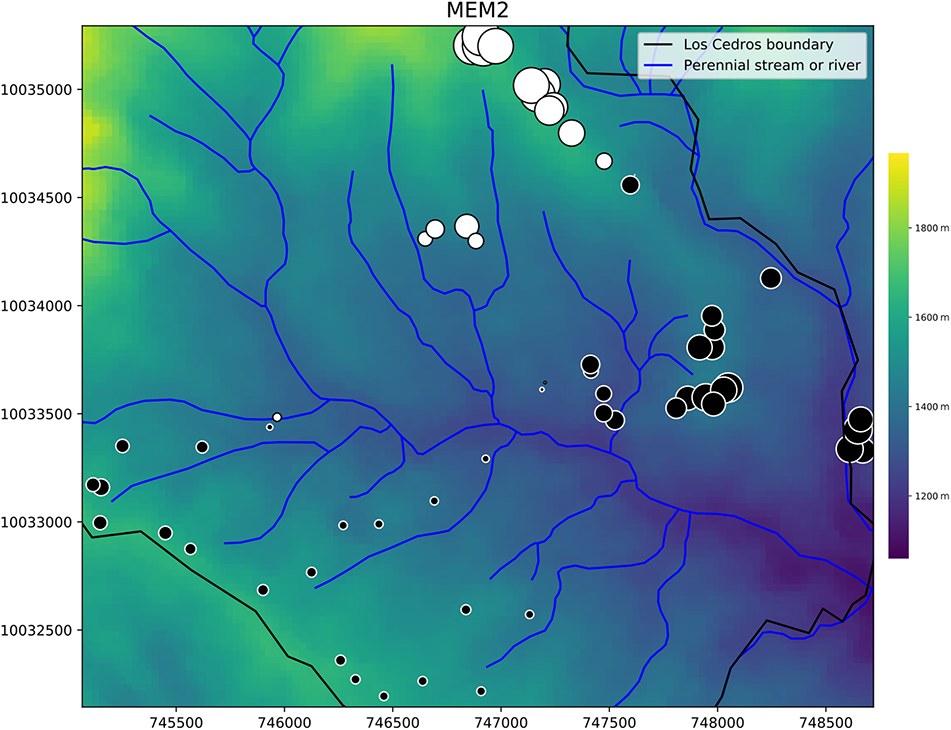

**Supplemental figure 9.**
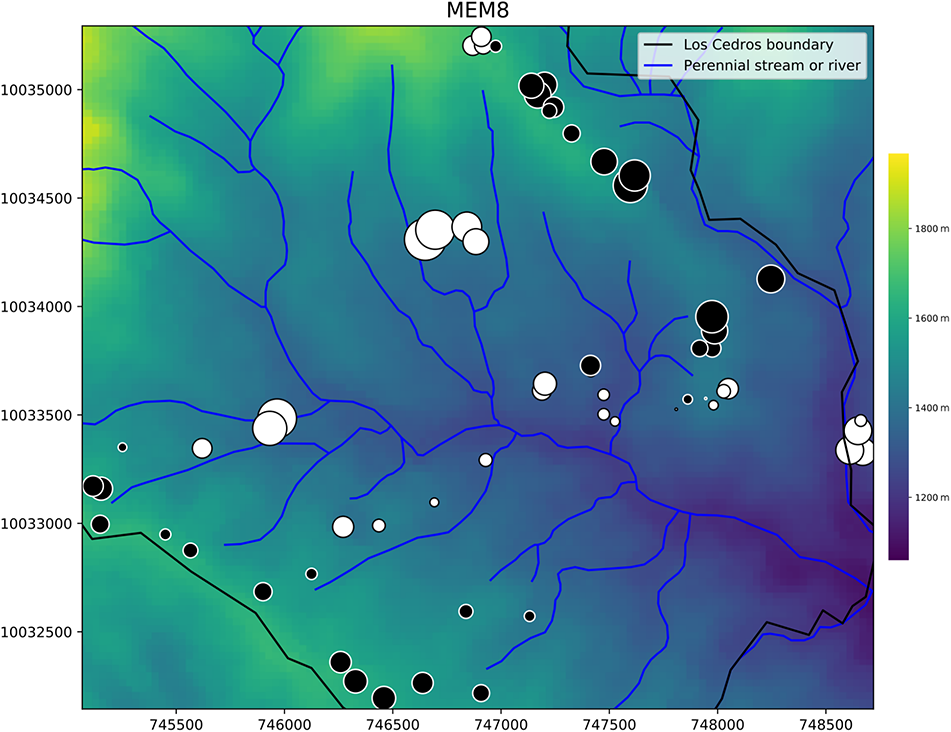

**Supplemental figure 10.**
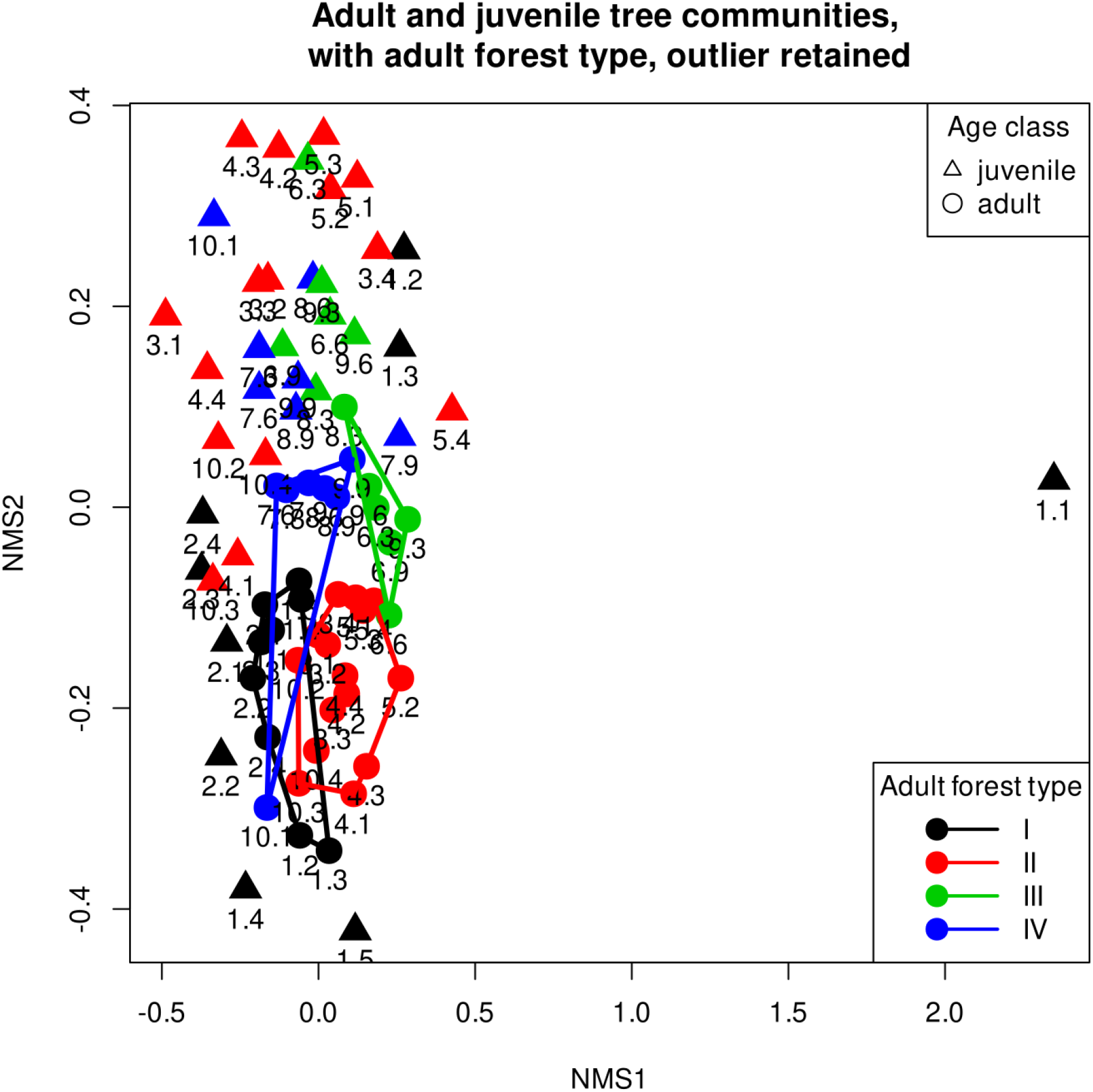

**Supplemental Table 1.**
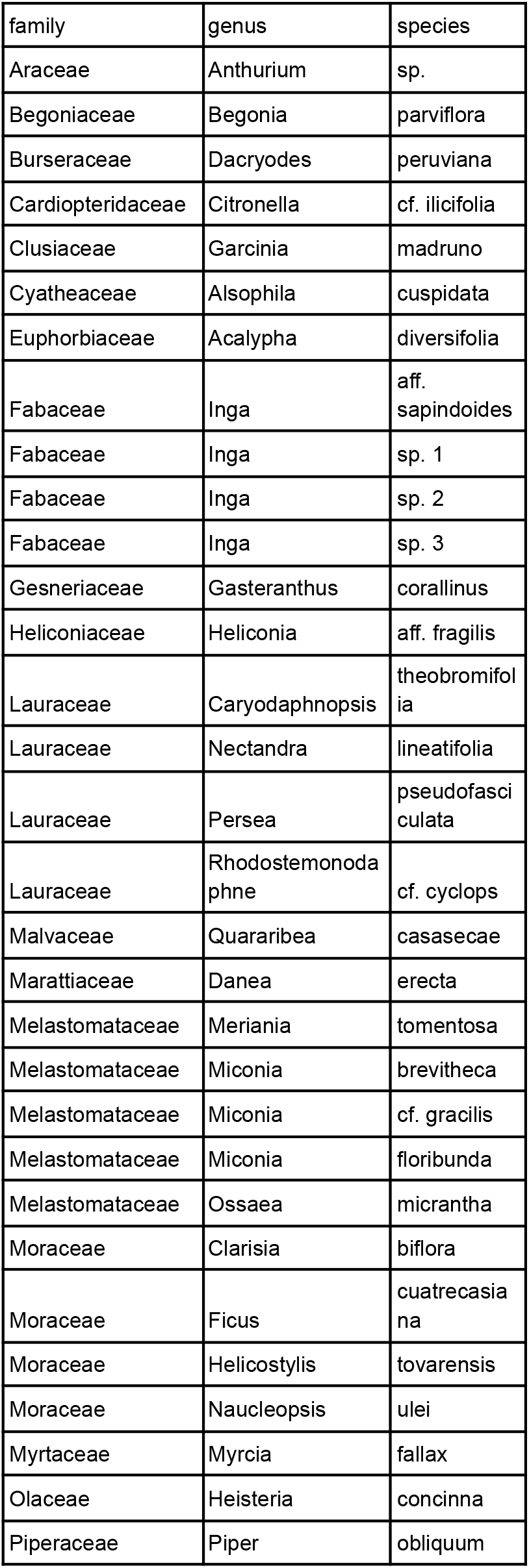
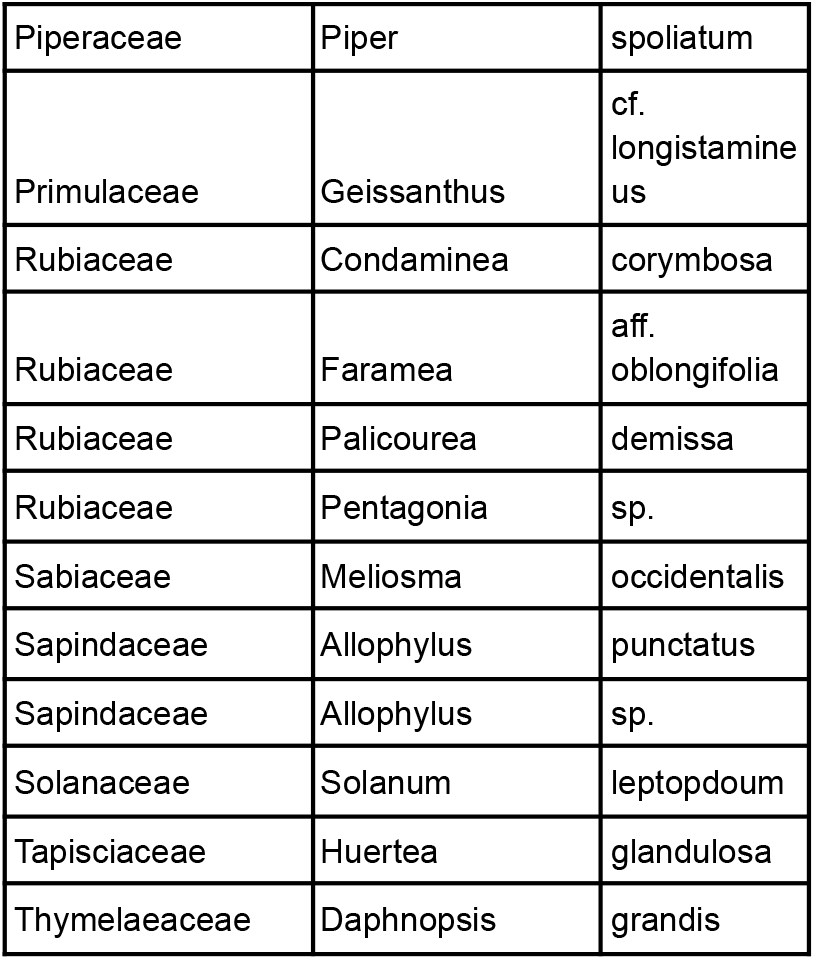
Species lists. **Supplemental Table 1A Permanent plot species list**

**Supplemental Table 1B.** Rapid survey species list

| family | genus | species |
| --- | --- | --- |
| Actinidaceae | Saurauia | sp. |
| Actinidaceae | sp. |  |
| Anacardiaceae | Mauria | cf. heterophylla |
| Anacardiaceae | Tapirira | guianensis |
| Annonaceae | Guatteria | cf. megalophylla |
| Annonaceae | Guatteria | megalophylla |
| Annonaceae | Guatteria | sp. |
| Annonaceae | Rollinia | aff. dolichopetala |
| Annonaceae | Rollinia | edulis |
| Annonaceae | sp. |  |
| Aquifoliaceae | Ilex | cf. yurumanguinis |
| Aquifoliaceae | Ilex | laurina |
| Araliaceae | Dendropanax | aff. macrophyllum |
| Araliaceae | Dendropanax | cf. macrophyllum |
| Araliaceae | Oreopanax | confusus |
| Araliaceae | Oreopanax | ecuadorensis |
| Arecaceae | Chamaedorea | pinnatifrons |
| Arecaceae | Hyospathe | elegans |
| Arecaceae | Wettinia | aff. oxycarpa |
| Arecaceae | Wettinia | quinaria |
| Asteraceae | Criptoniosis | cf. occidentalis |
| Asteraceae | Critoniopsis | occidentalis |
| Asteraceae | Piptocoma | discolor |
| Asteraceae | sp. |  |
| Asteraceae | Vernonanthura | patens |
| Bombacaceae | Matisia | aff. bolivarii |
| Bombacaceae | Matisia | castano |
| Bombacaceae | Matisia | sp. |
| Bombacaceae | Ochroma | pyramidale |
| Bombacaceae | Pachira | patinoi |
| Bombacaceae | Quararibea | aff. asterolepis |
| Bombacaceae | sp. |  |
| Boraginaceae | Cordia | aff. mexiana |
| Boraginaceae | Cordia | colombiana |
| Boraginaceae | Cordia | mexicana |
| Boraginaceae | Cordia | sp. |
| Boraginaceae | Tournefortia | cf. ramosissima |
| Brunelliaceae | Brunellia | acostae |
| Brunelliaceae | Brunellia | comocladifolia |
| Burseraceae | Dacryodes | cf. cupularis |
| Burseraceae | Dacryodes | cupularis |
| Burseraceae | Protium | ecuadorenses |
| Burseraceae | Protium | sp. |
| Burseraceae | sp. |  |
| Capparaceae | Capparis | macrophylla |
| Capparaceae | Capparis | sp. |
| Cecropiaceae | Cecropia | aff. andina |
| Cecropiaceae | Cecropia | aff. reticulata |
| Cecropiaceae | Cecropia | andina |
| Cecropiaceae | Cecropia | ficifolia |
| Cecropiaceae | Cecropia | reticulata |
| Cecropiaceae | Cecropia | sp.1 |
| Cecropiaceae | Cecropia | sp.2 |
| Cecropiaceae | Coussapoa | contorta |
| Chloranthaceae | Hedyosmum | cuatrecazanum |
| Chrysobalanaceae | Couepia | sp. |
| Chrysobalanaceae | Licania | durifolia |
| Chrysobalanaceae | Licania | sp. |
| Clusiaceae | Chrysochlamys | aff. dependens |
| Clusiaceae | Chrysochlamys | dependens |
| Clusiaceae | Chrysochlamys | sp. |
| Clusiaceae | Clusia | aff. dixonii |
| Clusiaceae | Clusia | aff. hydrogera |
| Clusiaceae | Clusia | dixonii |
| Clusiaceae | Clusia | sp. |
| Clusiaceae | Clusia | trochimorfis |
| Clusiaceae | Garcinia | macrophylla |
| Clusiaceae | Vismia | lauriformis |
| Cunoniaceae | Weinmannia | balbisiana |
| Cyatheaceae | Alsophila | cuspidata |
| Cyatheaceae | Alsophila | erinacea |
| Cyatheaceae | Cyathea | caracasana |
| Cyatheaceae | Cyathea | halonata |
| Cyatheaceae | sp. |  |
| Dichapetalaceae | Stephanopodium | angulatum |
| Dichapetalaceae | Stephanopodium | longipedicellatum |
| Dicksoniaceae | Dicksonia | sellowiana |
| Elaeocarpaceae | Sloanea | cf. grandiflora |
| Elaeocarpaceae | Sloanea | sp. 1 |
| Elaeocarpaceae | Sloanea | sp. 2 |
| Ericaceae | Psammisia | aberrans |
| Ericaceae | Psammisia | sodiroi |
| Erythroxylaceae | Erythroxylum | cf. novogranatense |
| Euphorbiaceae | Acalypha | aff. diversifolia |
| Euphorbiaceae | Acalypha | diversifolia |
| Euphorbiaceae | Alchornea | cf. grandis |
| Euphorbiaceae | Alchornea | cf. leptogyna |
| Euphorbiaceae | Hieronyma | alchorneoides |
| Euphorbiaceae | Hieronyma | asperifolia |
| Euphorbiaceae | Plukenetia | volubilis |
| Euphorbiaceae | Sapium | marmieri |
| Euphorbiaceae | Sapium | stylare |
| Euphorbiaceae | Tetrorchidium | andinum |
| Euphorbiaceae | Tetrorchidium | macrophyllum |
| Fab-Caesalpinieae | Senna | dariensis |
| Fab-Faboideae | Dussia | lehmannii |
| Fab-Faboideae | Erythrina | megistophylla |
| Fab-Faboideae | Erythrina | schimpffii |
| Fab-Faboideae | Erythrina | sp. |
| Fab-Faboideae | Lonchocarpus | sp. |
| Fab-Faboideae | sp. |  |
| Fab-Mimosoideae | Abarema | sp. |
| Fab-Mimosoideae | Cojoba | arborea |
| Fab-Mimosoideae | Inga | aff. lallensis |
| Fab-Mimosoideae | Inga | aff. nobilis |
| Fab-Mimosoideae | Inga | aff. sapindoides |
| Fab-Mimosoideae | Inga | aff. villosissima |
| Fab-Mimosoideae | Inga | lallensis |
| Fab-Mimosoideae | Inga | oerstediana |
| Fab-Mimosoideae | Inga | punctata |
| Fab-Mimosoideae | Inga | sp. |
| Fab-Mimosoideae | Inga | villosissima |
| Fab-Mimosoideae | sp. |  |
| Flacourtiaceae | Banara | guianensis |
| Flacourtiaceae | Banara | regia |
| Flacourtiaceae | Banara | riparia |
| Flacourtiaceae | Casearia | cajambrensis |
| Flacourtiaceae | Casearia | sylvestris |
| Flacourtiaceae | sp. |  |
| Hippocrataceae | Salacia | cordata |
| Indeterminado | sp. 1 |  |
| Indeterminado | sp. 10 |  |
| Indeterminado | sp. 11 |  |
| Indeterminado | sp. 12 |  |
| Indeterminado | sp. 13 |  |
| Indeterminado | sp. 14 |  |
| Indeterminado | sp. 15 |  |
| Indeterminado | sp. 16 |  |
| Indeterminado | sp. 17 |  |
| Indeterminado | sp. 18 |  |
| Indeterminado | sp. 19 |  |
| Indeterminado | sp. 2 |  |
| Indeterminado | sp. 3 |  |
| Indeterminado | sp. 4 |  |
| Indeterminado | sp. 5 |  |
| Indeterminado | sp. 6 |  |
| Indeterminado | sp. 7 |  |
| Indeterminado | sp. 8 |  |
| Indeterminado | sp. 9 |  |
| Lauraceae | Aniba | aff. hostmanniana |
| Lauraceae | Aniba | cf. hostmanniana |
| Lauraceae | Beilschmiedia | aff. costaricensis |
| Lauraceae | Beilschmiedia | alloiophylla |
| Lauraceae | Beilschmiedia | sp. |
| Lauraceae | Caryodaphnopsis | theobromifolia |
| Lauraceae | Cinnamomum | triplinerve |
| Lauraceae | Endlicheria | cf. chalisea |
| Lauraceae | Endlicheria | cf. sericea |
| Lauraceae | Endlicheria | cf. tessmannii |
| Lauraceae | Endlicheria | formosa |
| Lauraceae | Endlicheria | klugii |
| Lauraceae | Endlicheria | sp. |
| Lauraceae | Licaria | aff. tomentosa |
| Lauraceae | Licaria | cf. tomentosa |
| Lauraceae | Nectandra | acutifolia |
| Lauraceae | Nectandra | aff. guararipo |
| Lauraceae | Nectandra | aff. laurel |
| Lauraceae | Nectandra | aff. membranacea |
| Lauraceae | Nectandra | aff. parviflora |
| Lauraceae | Nectandra | aff. Purpurea |
| Lauraceae | Nectandra | guararipo |
| Lauraceae | Nectandra | membranacea |
| Lauraceae | Nectandra | obtusata |
| Lauraceae | Nectandra | sp. 1 |
| Lauraceae | Nectandra | sp. 2 |
| Lauraceae | Nectandra | sp. 3 |
| Lauraceae | Nectandra | sp.1 |
| Lauraceae | Ocotea | aff. cernua |
| Lauraceae | Ocotea | aff. floccifera |
| Lauraceae | Ocotea | aff. heterochroma |
| Lauraceae | Ocotea | aff. insularis |
| Lauraceae | Ocotea | aff. oblonga |
| Lauraceae | Ocotea | cernua |
| Lauraceae | Ocotea | floccifera |
| Lauraceae | Ocotea | insularis |
| Lauraceae | Ocotea | macrophylla |
| Lauraceae | Ocotea | oblonga |
| Lauraceae | Ocotea | sp. 1 |
| Lauraceae | Ocotea | sp. 2 |
| Lauraceae | Ocotea | stenoneura |
| Lauraceae | Persea | aff. areolatocostae |
| Lauraceae | Persea | aff. pseudofasciculata |
| Lauraceae | Persea | povedae |
| Lauraceae | Persea | pseudofasciculata |
| Lauraceae | Persea | rigens |
| Lauraceae | Pleurothyrium | aff. giganthum |
| Lauraceae | Pleurothyrium | cinereum |
| Lauraceae | Pleurothyrium | giganthum |
| Lauraceae | Pleurothyrium | glabritepalum |
| Lauraceae | Pleurothyrium | sp. |
| Lauraceae | Rhodostemonoda<br>phne | aff. cyclops |
| Lauraceae | Rhodostemonoda<br>phne | cyclops |
| Lauraceae | sp. 1 |  |
| Lauraceae | sp. 2 |  |
| Lauraceae | sp. 3 |  |
| Lauraceae | sp. 4 |  |
| Lecythidaceae | Eschweilera | caudiculata |
| Lecythidaceae | Eschweilera | integrifolia |
| Lecythidaceae | Eschweilera | rimbachii |
| Lecythidaceae | Gustavia | dodsonii |
| Magnoliaceae | Talauma | cf. gilbertoi |
| Malphigiaceae | sp. |  |
| Malvaceae | Wercklea | ferox |
| Malvaceae | Wercklea | sp. |
| Melastomataceae | Conostegia | aff. centronioides |
| Melastomataceae | Conostegia | cuatrecasii |
| Melastomataceae | Conostegia | superba |
| Melastomataceae | Leandra | subseriata |
| Melastomataceae | Meriania | finicola |
| Melastomataceae | Meriania | maxima |
| Melastomataceae | Meriania | tomentosa |
| Melastomataceae | Miconia | aff. brevitheca |
| Melastomataceae | Miconia | aggregata |
| Melastomataceae | Miconia | cf. subnodosa |
| Melastomataceae | Miconia | dapsiliflora |
| Melastomataceae | Miconia | loveyoides |
| Melastomataceae | Miconia | sp. |
| Melastomataceae | Ossaea | cf. micrantha |
| Melastomataceae | Ossaea | micrantha |
| Melastomataceae | Ossaea | robusta |
| Melastomataceae | sp. 1 |  |
| Melastomataceae | sp. 3 |  |
| Melastomataceae | sp.1 |  |
| Melastomataceae | sp.2 |  |
| Melastomataceae | sp.3 |  |
| Melastomataceae | Topobea | dodsonorum |
| Meliaceae | Guarea | aff. macrophylla |
| Meliaceae | Guarea | cartaguenya |
| Meliaceae | Guarea | cf. pubescens |
| Meliaceae | Guarea | cf. subandina |
| Meliaceae | Guarea | kunthiana |
| Meliaceae | Ruagea | aff. glabra |
| Meliaceae | Ruagea | cf. glabra |
| Meliaceae | Ruagea | cf. pubescens |
| Meliaceae | Ruagea | glabra |
| Meliaceae | Ruagea | pubescens |
| Meliaceae | Ruagea | trisperma |
| Meliaceae | sp. 1 |  |
| Meliaceae | sp.1 |  |
| Meliaceae | sp.2 |  |
| Meliaceae | Trichilia | aff. septentrionalis |
| Meliaceae | Trichilia | septentrionalis |
| Meliaceae | Trichilia | sp. |
| Meliaceae | Trichilia | sp. 1 |
| Monimiaceae | Mollinedia | sp. |
| Moraceae | Clarisia | biflora |
| Moraceae | Ficus | andicola |
| Moraceae | Ficus | caldasiana |
| Moraceae | Ficus | cf. caldasiana |
| Moraceae | Ficus | cotopaxiensis |
| Moraceae | Ficus | cuatrecasana |
| Moraceae | Ficus | dulciaria |
| Moraceae | Ficus | macbridei |
| Moraceae | Ficus | maxima |
| Moraceae | Ficus | mutisii |
| Moraceae | Ficus | sibundoya |
| Moraceae | Ficus | sp. |
| Moraceae | Ficus | subandina |
| Moraceae | Ficus | tonduzii |
| Moraceae | Ficus | trianae |
| Moraceae | Helicostylis | tomentosa |
| Moraceae | Helicostylis | tovarensis |
| Moraceae | Morus | insignis |
| Moraceae | Naucleopsis | naga |
| Moraceae | Perebea | aff. xanthochyma |
| Moraceae | Pseudolmedia | macrophylla |
| Moraceae | Pseudolmedia | rigida |
| Moraceae | Sorocea | jaramilloi |
| Moraceae | Sorocea | steinbachii |
| Moraceae | sp. |  |
| Moraceae | sp. 1 |  |
| Moraceae | Trophis | racemosa |
| Moraceae | vs. Perebea | angustifolia |
| Myristicaceae | Otoba | gordoniiifolia |
| Myristicaceae | Virola | aff. sebifera |
| Myristicaceae | Virola | sp. |
| Myrsinaceae | Ardisia | cf. colombiana |
| Myrsinaceae | Cybianthus | humilis |
| Myrsinaceae | Geissanthus | aff. longistamineus |
| Myrsinaceae | Geissanthus | aff. pinchinchana |
| Myrsinaceae | Stylogyne | ambigua |
| Myrsinaceae | Weigeltia | cf. goudotiana |
| Myrtaceae | Calyptranthes | sp. |
| Myrtaceae | Eugenia | cf. florida |
| Myrtaceae | Eugenia | florida |
| Myrtaceae | Eugenia | sp. |
| Myrtaceae | Myrcia | aff. aliena |
| Myrtaceae | Myrcia | aff. fallax |
| Myrtaceae | Myrcia | cf. aliena |
| Myrtaceae | Myrcia | cf. fallax |
| Myrtaceae | Myrcia | fallax |
| Myrtaceae | Myrcia | splendens |
| Myrtaceae | sp. |  |
| Ochnaceae | Ouratea | aff. weberbaueri |
| Ochnaceae | Ouratea | weberbaueri |
| Olacaceae | Heisteria | sp. |
| Piperaceae | Piper | aduncum |
| Piperaceae | Piper | aff. nudibracteatum |
| Piperaceae | Piper | cf. imperiale |
| Piperaceae | Piper | fuliginosum |
| Piperaceae | Piper | imperiale |
| Piperaceae | Piper | sp. |
| Rubiaceae | Agouticarpa | grandistipula |
| Rubiaceae | Condaminea | corymbosa |
| Rubiaceae | Elaeagia | cf. karstenii |
| Rubiaceae | Elaeagia | karstenii |
| Rubiaceae | Elaeagia | utilis |
| Rubiaceae | Faramea | aff. calyptrata |
| Rubiaceae | Faramea | flavicans |
| Rubiaceae | Faramea | fragans |
| Rubiaceae | Faramea | glandulosa |
| Rubiaceae | Faramea | oblongifolia |
| Rubiaceae | Faramea | sp. |
| Rubiaceae | Guettarda | crispiflora |
| Rubiaceae | Guettarda | hirsuta |
| Rubiaceae | Hoffmannia | cf. parviflora |
| Rubiaceae | Isertia | laevis |
| Rubiaceae | Isertia | pittieri |
| Rubiaceae | Palicourea | acanthacea |
| Rubiaceae | Palicourea | aff. chignul |
| Rubiaceae | Palicourea | amethystina |
| Rubiaceae | Palicourea | cf. stipularis |
| Rubiaceae | Palicourea | chimboracensis |
| Rubiaceae | Palicourea | gentryi |
| Rubiaceae | Palicourea | gibbosa |
| Rubiaceae | Palicourea | sp. |
| Rubiaceae | Palicourea | stipularis |
| Rubiaceae | Palicourea | thyrsiflora |
| Rubiaceae | Pentagonia | macrophylla |
| Rubiaceae | Psychotria | cf. steyermarkii |
| Rubiaceae | Psychotria | flaviflora |
| Rubiaceae | Psychotria | gentryi |
| Rubiaceae | Psychotria | paeonia |
| Rubiaceae | Psychotria | pilosa |
| Rubiaceae | Rondeletia | reflexa |
| Rubiaceae | sp. 1 |  |
| Rubiaceae | sp.1 |  |
| Rubiaceae | sp.2 |  |
| Rubiaceae | sp.3 |  |
| Rubiaceae | Tocoyena | pittieri |
| Rutaceae | Zanthoxylum | mauriifolium |
| Rutaceae | Zanthoxylum | sp. |
| Sabiaceae | Meliosma | aff. occidentalis |
| Sabiaceae | Meliosma | occidentalis |
| Sapindaceae | Allophyllus | floribundus |
| Sapindaceae | Allophyllus | incanus |
| Sapindaceae | Allophyllus | sp. |
| Sapindaceae | Cupania | aff. cinerea |
| Sapindaceae | Talisia | aff. nervosa |
| Sapindaceae | Talisia | cf. nervosa |
| Sapotaceae | Chrysophyllum | aff. venezuelanense |
| Sapotaceae | Chrysophyllum | cf. argenteum |
| Sapotaceae | Chrysophyllum | manaosense |
| Sapotaceae | Chrysophyllum | sp. |
| Sapotaceae | Pouteria | torta |
| Simaroubaceae | Simarouba | amara |
| Siparumaceae | Siparuna | laurifolia |
| Siparumaceae | Siparuna | salvioides |
| Siparumaceae | Siparuna | sp. |
| Siparunaceae | Siparuna | salvioides |
| Solanaceae | Cestrum | megalophyllum |
| Solanaceae | Markea | pavonii |
| Solanaceae | Solanum | aff. circinatum |
| Solanaceae | Solanum | aff. cucullatum |
| Solanaceae | Solanum | cf. circinatum |
| Solanaceae | Solanum | cucullatum |
| Solanaceae | Solanum | lepidotum |
| Solanaceae | Solanum | sp. |
| Staphyleaceae | Huetea | glandulosa |
| Staphyleaceae | Turpinia | occidentalis |
| Styracaceae | Styrax | weberbaueri |
| Symplocaceae | Symplocos | fuliginosa |
| Theaceae | Freziera | reticulata |
| Theophrastaceae | Clavija | eggersiana |
| Theophrastaceae | Clavija | membranacea |
| Thymelaeaceae | Schoenobiblus | cf. daphnoides |
| Thymelaeaceae | Schoenobiblus | daphnoides |
| Tiliaceae | sp. |  |
| Ulmaceae | Trema | micrantha |
| Urticaceae | Boehmeria | caudata |
| Urticaceae | Phenax | aff. rugosus |
| Urticaceae | Urera | aff. caracasana |
| Urticaceae | Urera | baccifera |
| Urticaceae | Urera | caracasana |
| Verbenaceae | Aegiphila | alba |
| Violaceae | Gloeospermum | longifolium |

**Supplemental Table 2.**
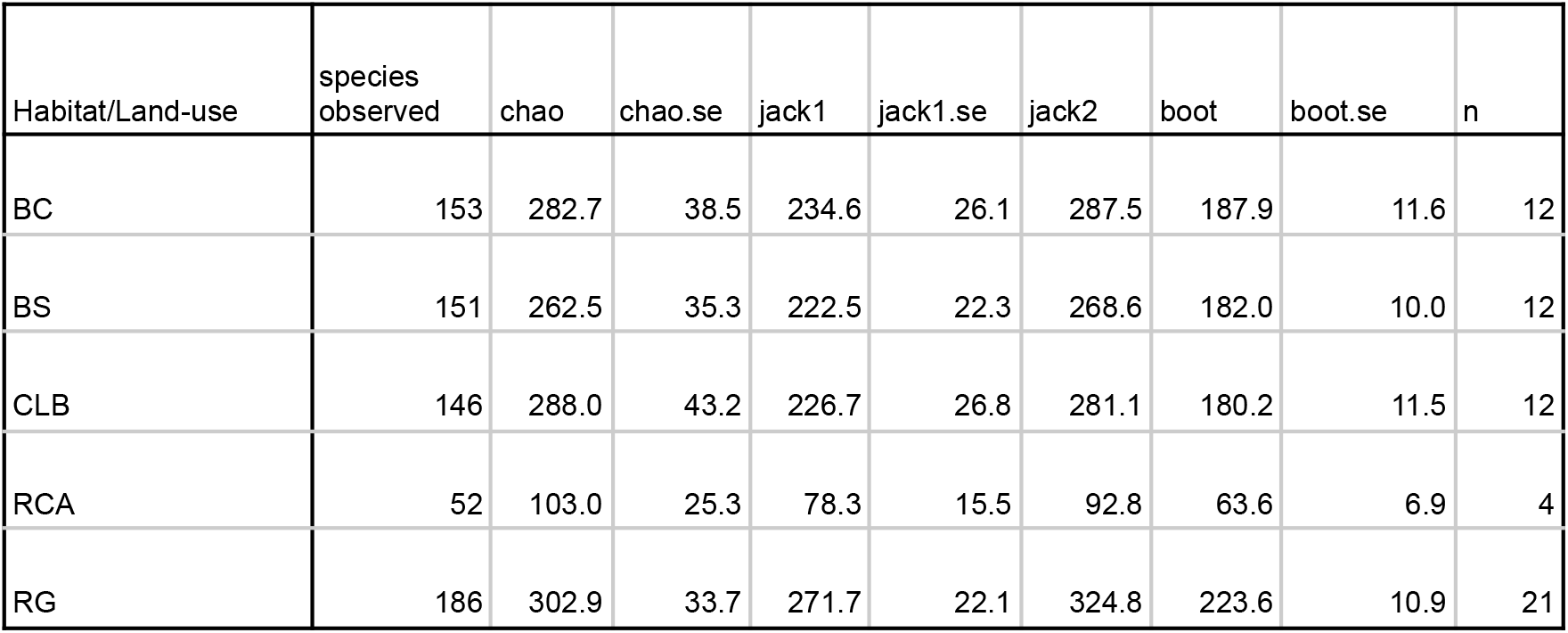
Species diversity estimates for historical land-use/habitat types

| Habitat/Land-use | species<br>observed | chao | chao.se | jack1 | jack1.se | jack2 | boot | boot.se | n |
| --- | --- | --- | --- | --- | --- | --- | --- | --- | --- |
| BC | 153 | 282.7 | 38.5 | 234.6 | 26.1 | 287.5 | 187.9 | 11.6 | 12 |
| BS | 151 | 262.5 | 35.3 | 222.5 | 22.3 | 268.6 | 182.0 | 10.0 | 12 |
| CLB | 146 | 288.0 | 43.2 | 226.7 | 26.8 | 281.1 | 180.2 | 11.5 | 12 |
| RCA | 52 | 103.0 | 25.3 | 78.3 | 15.5 | 92.8 | 63.6 | 6.9 | 4 |
| RG | 186 | 302.9 | 33.7 | 271.7 | 22.1 | 324.8 | 223.6 | 10.9 | 21 |

